# Peri-Head Distance Coding in the Mouse Brainstem

**DOI:** 10.64898/2025.11.30.691334

**Authors:** Wenxi Xiao, Kyle S. Severson, Hao Zheng, Ke Chen, P. M. Thompson, Manuel S. Levy, Seonmi Choi, Shengli Zhao, Jun Takatoh, Vincent Prevosto, Fan Wang

**Author notes:** These two authors contributed equally (Wenxi Xiao, Kyle S. Severson).

## Abstract

Perceiving object distance in peri-personal space is essential for guiding movement and avoiding danger. During active sensation, distance information is often anchored to the body via touch; yet how early somatosensory circuits extract distance information from tactile inputs remains unclear. Here, we investigate how second-order neurons in the mouse whisker brainstem encode peri-head distance. Using *in vivo* extracellular recordings in awake mice in a naturalistic wall-passing paradigm, we find brainstem neurons employ two distance-coding schemes: a “proximity” code, where firing increases monotonically as objects approach the face; and a “map” code, where neurons exhibit peak tuning at specific distances to collectively tile peri-head space. The map code outperforms proximity code in population decoding of distance. Perturbation experiments reveal multi-whisker integration and internuclear inhibition contribute to the generation of map-like tuning. These findings highlight a previously underappreciated computational role for brainstem circuits, where inhibition acts as a neural comparator to transform proximity-based sensory inputs into a map-like representation of peri-personal space.

## MAIN TEXT

The ability to perceive and represent space is fundamental for animals to interact with the external world. Estimating the distance of a predator, planning reach-to-grasp movements for food, or navigating complex terrains with obstacles and gaps all require accurate spatial processing. Of particular ethological importance is the space immediately surrounding the body, peripersonal space (PPS), which serves as a critical interface for sensorimotor integration, enabling object interaction, social engagement, and defensive behaviors (Serino, 2019). Classic primate studies identified multisensory neural substrates for PPS, where bimodal neurons respond both to tactile stimulation on specific body parts and to visual or auditory stimuli presented *near* those same body surfaces (Graziano & Cooke, 2006).

PPS has been proposed as a contact-related action field (Bufacchi & Iannetti, 2018), in which an accurate representation of relative distance to nearby entities directly constrains motor planning. For example, reaching for a cup beside the hand versus one placed farther away requires distinct patterns of arm and trunk muscle activation; likewise, crossing a narrow versus wide gap imposes different mechanical and postural demands. Some PPS-encoding neurons increase their firing as visual stimuli move into near space and loom toward the body surface, consistent with a proximity-based rate code (Graziano & Cooke, 2006). However, neuropsychological evidence indicates that distance from the body—in effect, what lies within PPS—is a fundamental component of spatial representation (Cowey et al., 1994; Làdavas et al., 1998), suggesting that a simple monotonic proximity code may be insufficient as the sole format for encoding distances in PPS. Clarifying how distance is represented in PPS is therefore crucial for uncovering the sensory computations that support spatial awareness and action execution.

To estimate distances in the surrounding environment, humans often rely on physical references—most commonly, the body itself—for example, describing something as being “an arm’s length away”. This body-referenced approach to spatial perception dates back to ancient civilizations, which formalized units of measurement based on body parts, including the “palm”, “foot”, “step”, and “fathom” (the span of outstretched arms). Compared to continuous proximity cues, body-part references provide discrete spatial scales that can be more readily interpreted and used for motor planning. This intuitive measurement strategy raises the question of whether the brain implements an analogous “internal ruler” that systematically encodes distances relative to specific body parts. If so, what is the representational format of this distance code? Addressing these questions may provide important insights into how spatial representations within PPS are constructed and used by the brain to guide behavior.

The rodent whisker sensory system provides a natural framework to study distance encoding, with grid-like whiskers serving as a built-in coordinate system for peri-head space (Brecht et al., 1997). Rodents are nocturnal animals that rely extensively on their whiskers to explore their surroundings (Diamond et al., 2008). The whisker array is organized into arcs whose lengths form a gradient along the anterior-posterior axis (Belli et al., 2017), mapping whisker identity onto specific regions of peri-head space (Figure 1A). Each whisker thus serves as a potential distance reference, allowing the brain to integrate inputs across the array into a head-centric representation of peri-head space (Pluta et al., 2017). Supporting this idea, a subset of neurons in the whisker barrel cortex (wS1) exhibit non-monotonic tuning to specific distances between the mouse’s head and surrounding walls during virtual corridor navigation (Sofroniew & Vlasov et al., 2015). Such tuning can be interpreted as a “map code”, where the peaks of the tuning curves map the distance to the head like tick marks on a ruler. At the population level, these preferred distances spanned the full range of whisker reach, effectively forming a head-centered representation of peri-head distance.

**Figure 1.**
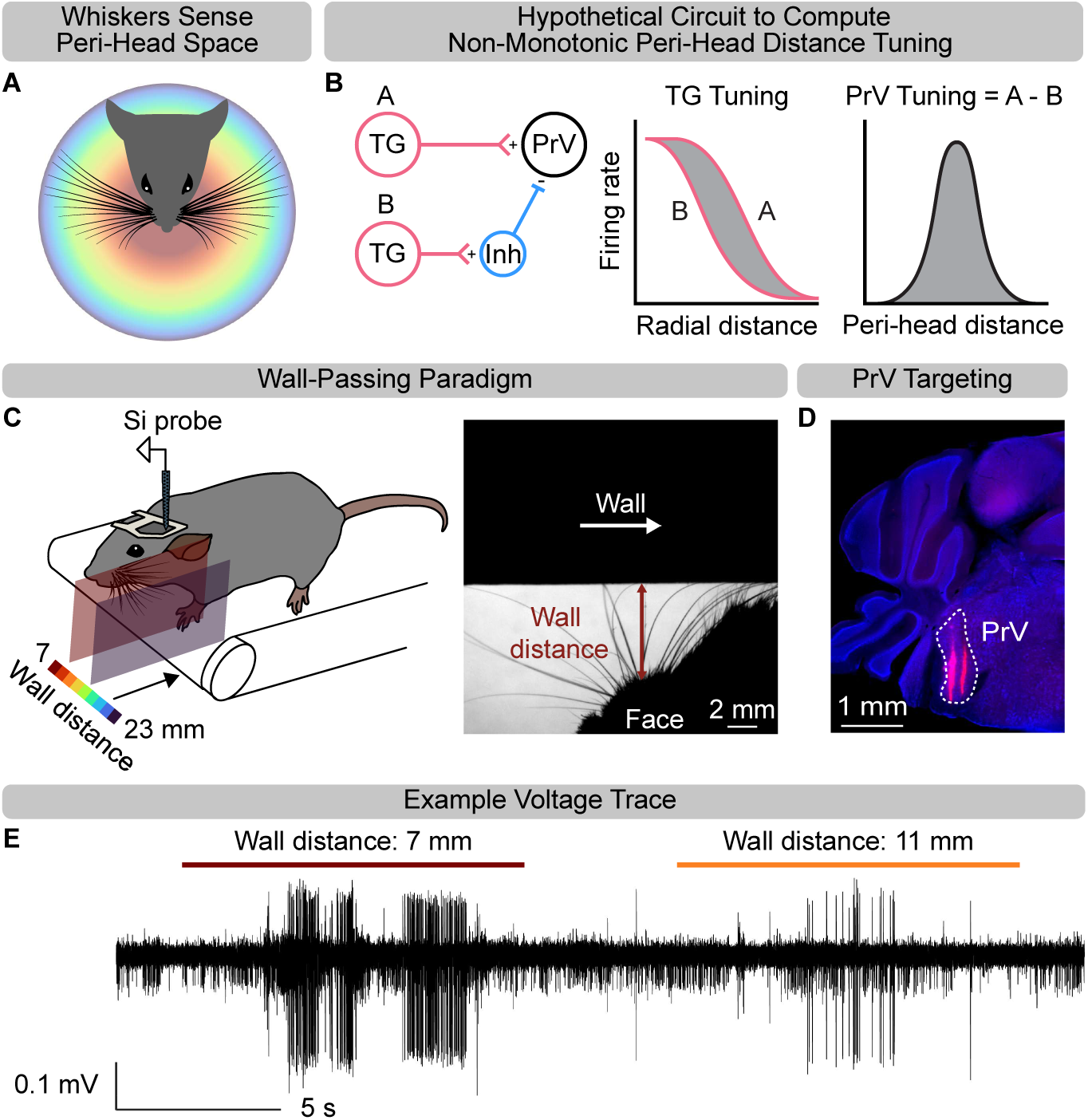
Acute PrV recordings in a wall passing paradigm. (**A**) Conceptual illustration showing how whiskers map peri-head space in head-centered coordinates. (**B**) Left, schematic of a hypothetical brainstem circuit to compute non-monotonic distance code. Two TG neurons (A and B, red) with different distance sensitivities project to the brainstem: neuron A provides excitatory input to a PrV output neuron (C, black), whereas neuron B synapses onto an inhibitory neuron (blue) in the trigeminal brainstem complex. Middle, tuning curves for neurons A and B; the shaded region indicates the difference between their distance-tuning profiles. Right, linear summation yields a non-monotonic tuning curve in neuron C, with a peak at an intermediate peri-head distance. (**C**) Left, Schematic of the experimental setup. A head-fixed mouse ran on a treadmill while a wall moved from anterior to posterior past the mouse’s face. Wall distance from the face was randomly varied between 7-23 mm in 2-mm increments on each pass. Each recording contains 15-20 repetitions per distance. Right, Example video frame showing multiple whiskers contacting the wall (moving left to right). Red arrow indicates measured distance between face and wall. (**D**) Example coronal brain section (4.8 mm posterior to Bregma) showing two silicon probe tracks through PrV labeled with Dil (red); cell bodies stained with Nissl (blue). (**E**) Example extracellular voltage trace from one recording session. Colored bars indicate wall-passing periods.

However, the map code for peri-head distance has not been observed in the periphery. Whiskers are innervated by mechanosensory neurons with soma residing in the trigeminal ganglion (TG), each of which has a single-whisker receptive field (Petersen, 2007). These neurons respond to mechanical stresses on the whisker follicle generated when the whisker bends upon contact with an object (Mitchinson et al., 2004; Bush et al., 2016; Campagner et al., 2016; Severson & Xu et al., 2017). Consequently, extracellular recordings from TG neurons, using a vertical pole positioned at various locations along the length of a single whisker, revealed that firing rates increase as the point of contact moves closer to the face (Szwed et al., 2006). Thus, in the periphery, TG neurons encode radial distance via a proximity-based rate code, referenced to the base of a single whisker (Ahissar & Knutsen, 2008). This implies that a sensory code transformation occurs along the whisker sensory pathway to convert whisker-centered proximity code in the periphery into the map-like peri-head distance code observed in wS1.

We hypothesize that a transformation from proximity to map-like representation of peri-head distance occurs at the level of second-order neurons in the principal trigeminal nucleus (PrV) of the brainstem. PrV contains neurons with either single- or multi-whisker receptive fields (Minnery & Simons, 2003) and receives convergent inputs from multiple subtypes of TG mechanosensory neurons, including both rapidly and slowly adapting afferents (Sakurai et al., 2013; Jacquin et al., 2015). In addition, PrV circuits incorporate local glycinergic inhibitory interneurons (Avendaño et al., 2005; Bellavance et al., 2010) and receive topographically organized long-range GABAergic input from the spinal trigeminal nucleus interpolaris (SpVi; Jacquin et al., 2015, Furuta et al., 2008). These circuit motifs, supporting both multi-whisker integration (Minnery & Simons, 2003; Furuta et al., 2008) and feedforward inhibition, make PrV a candidate site for implementing the transformation of whisker-centered proximity signals into a head-centered map code. Inspired by Porter and Groh (2006), we propose that a simple circuit (Figure 1B) could transform monotonic proximity-based inputs from TG neurons into a non-monotonic map-like readout of peri-head distance. Importantly, this circuit requires inhibition: when proximity-based inputs from two TG neurons with different mechanical sensitivity are subtracted via inhibition, a peak response at a specific peri-head distance emerges. To test these hypotheses, we performed *in vivo* electrophysiological recordings from PrV neurons in a wall-passing paradigm.

## Results

### A wall-passing paradigm to study peri-head distance encoding in the whisker brainstem

To evoke tactile sensation similar to that experienced during naturalistic wall-following behavior, we devised a wall-passing paradigm for awake, behaving mice head-fixed on a linear treadmill (Severson & Xu et al., 2017; Figure 1C, left; STAR Methods). In this setup, each stimulus presentation consists of a wall moving past the mouse’s head at a fixed radial distance from anterior to posterior, delivering controlled tactile stimulation to the whisker array. A linear actuator moved the wall along the anteroposterior axis at a constant velocity of 10 mm/s, operating independently of the animal’s locomotion (open loop). Before each wall pass, the radial distance between the wall and the mouse’s face (wall distance) was randomly set. Each pass lasted approximately 14 seconds, with wall movement onset at t = 0 and whisker-wall contacts typically occurring between 3-11 seconds. In total, each distance was presented for 15-20 wall stimulus presentations. The wall position was aligned to the midline of the head, with wall distances measured from the wall to the C2 whisker follicle (Figure 1C, right).

After habituation to the wall-passing stimulus, mice exhibited predominantly passive whisker-wall interactions. To assess the degree of active retraction, we compared wall-induced whisker bending angles of a single row of whiskers in awake versus anesthetized conditions (see Supplemental Figure S1). Under anesthesia, the moving wall pushed whiskers in the retraction direction, producing larger absolute bending angles at closer wall distances (Supplemental Figure S1D, right). Awake, behaving mice in the same wall-passing paradigm showed similar whisker bending angles (mean difference: 2.28°), with no statistically significant difference from the anesthetized condition across whiskers (Supplemental Figure S1F). This observation suggests that mice engaged in little active whisker retraction to avoid contact, instead allowing the whiskers to passively glide along the passing wall. Given this minimal wall-induced active retraction, whisker bending mostly increased monotonically with wall proximity, supporting proximity-based distance encoding in the periphery.

### PrV contains both proximity and map codes for peri-head distance

To examine how PrV neurons respond to the passing wall stimulus at varying distances, we first performed *in vivo* extracellular recordings in mice with only the C-row whiskers intact, using single tungsten electrodes (see Supplemental Figure S2A-C; STAR Methods). Based on our single-unit isolation metrics (see STAR Methods; Supplemental Figure S3J-M), we isolated 46 units from 14 mice (see Table S1 for mouse metadata), including 18 units with single-whisker and 28 units with multi-whisker receptive fields. We analyzed single-unit responses using peri-event time histograms (PETHs) by computing the averaged firing rates for wall contact (3-11 s) and baseline periods (−4.5-0 s) across trials. To determine wall responsiveness, we applied one-tailed Wilcoxon signed-rank tests (see STAR Methods). Of the 46 units, 41/46 (89.1%) were wall-activated and 3/46 (6.5%) were wall-suppressed; one unit showed both activation and suppression at different distances, yielding a total of 43/46 (93.5%) wall-responsive (presumably touch-sensitive) units. To assess distance tuning, we performed two-tailed Wilcoxon signed-rank tests across all distance pairs (p < 0.05, corrected using the Benjamini-Hochberg false discovery rate). Units with significant differences in mean firing rates for at least one distance pair were classified as “distance-tuned” (84.8%, 39/46). For each distance-tuned unit, we computed a z-scored tuning curve (see STAR Methods). The majority of units (52.2%, 24/46) showed a broadly monotonic, inversely proportional relationship with wall distance (e.g., Supplemental Figure S2D), consistent with PrV neurons inheriting proximity-based TG inputs. Interestingly, a distinct subset of units (13.0%, 6/46) clearly preferred intermediate or far wall distances (e.g., Supplemental Figure S2E), revealing a heterogeneous representation of peri-head distance in PrV (see Supplemental Figure S2F).

Because high-impedance tungsten electrodes preferentially capture high-amplitude spikes, unit isolation was biased toward larger neurons (Buzsáki et al., 2012), raising the possibility that under-sampling biased our estimate of the proportion of cells tuned to intermediate wall distances. Furthermore, the low-yield of single electrode recordings resulted in a small sample size (n=46 units well-isolated units over 14 mice). To overcome these limitations, we shifted to 32-channel single-shank silicon probes for acute recordings of PrV single-unit activity during the wall-passing paradigm (Figure 1C-E; See STAR Methods). We isolated 1,141 single units across 36 mice (Supplemental Figure S3A-E). Of the 1,141 units, 67.3% (768/1141) were significantly modulated by the passing-wall stimulus (“wall-responsive”). Among these wall-responsive units, 532/768 (69.3%) showed significant activation at one or more distances, 333/768 (43.4%) showed significant suppression, and 97 units met both criteria (activated at some distances and suppressed at others). Given that our probe spanned the entire dorsoventral axis of PrV (Figure 1D) and that whisker-responsive neurons are specifically localized to ventral PrV (barrelettes, Bechara et al., 2015), the prominence of unresponsive units, likely corresponding to unstimulated microvibrissa-, jaw-, or lip-related neurons in dorsal PrV, is not unexpected.

Representative spike rasters, PETHs, and z-scored wall-distance tuning curves from six wall-responsive units are shown in Figure 2A-C. These examples illustrate the diversity of tuning responses: Unit 1 showed monotonic increases in firing with decreasing wall distance (“proximity” tuning); Units 2-4 exhibited peak firing at specific intermediate distances (“map” tuning); Unit 5 displayed a complex response profile with both suppression and activation depending on wall distance; and Unit 6 was suppressed during wall contact. In addition to sustained responses, wall-touch onset and offset responses were often observed in the same PrV neurons, suggesting convergence of multiple afferent types (i.e., slowly and rapidly adapting) onto PrV neurons, consistent with anatomical evidence that different types of TG afferent terminals are intermingled within PrV barrelettes (Sakurai et al., 2013). However, our analyses focused on mean firing rates across the entire wall-contact period (8 s) and did not further dissect these transient dynamics.

**Figure 2.**
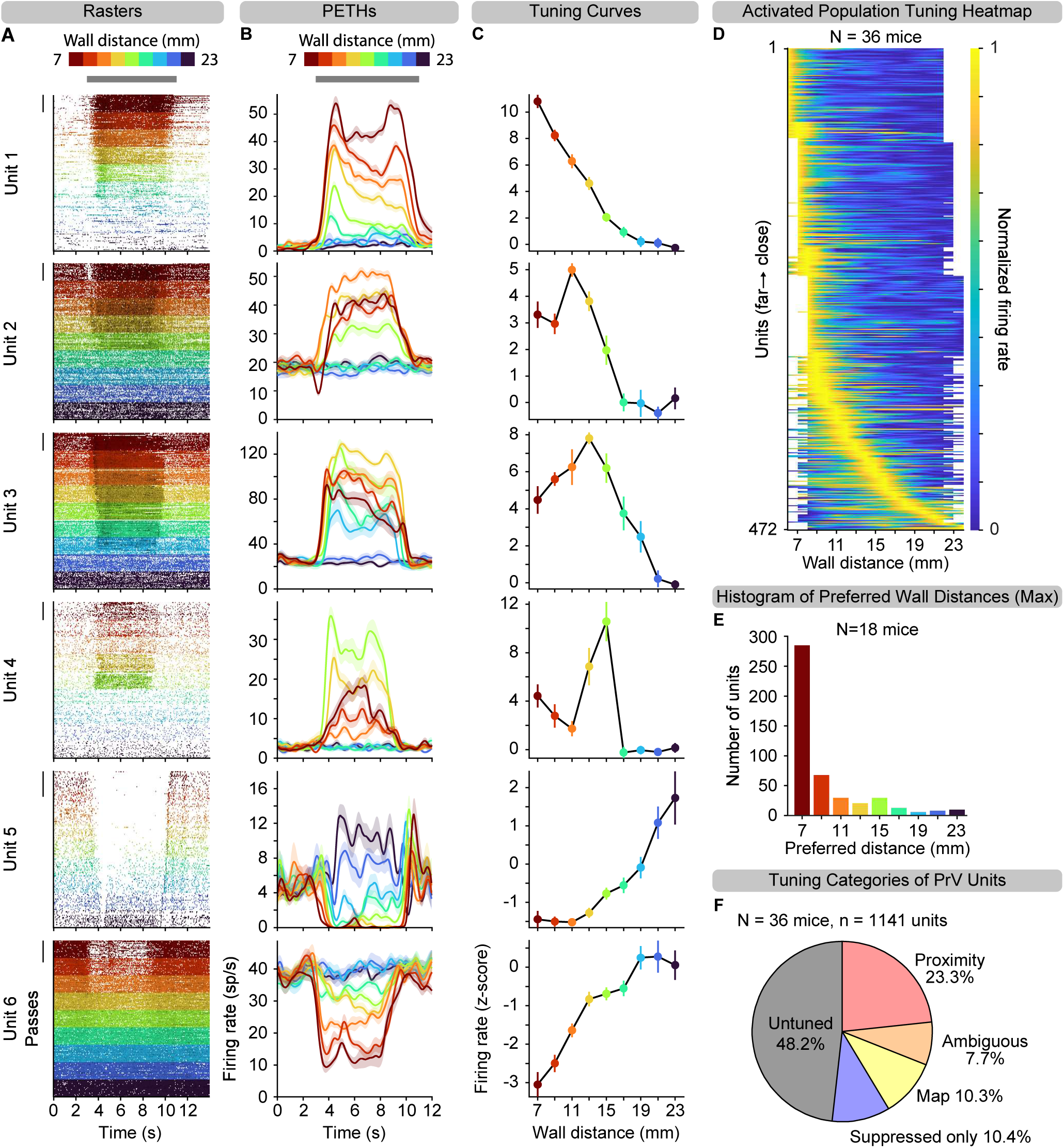
PrV contains multiple neural codes for peri-head distance. (**A**) Example spike rasters of six single units (units 1-6) in PrV during repeated wall pass trials. Each row corresponds to one wall pass trial, and each dot corresponds to a time bin (50 ms) containing at least one spike, colored according to wall distances. Wall passing begins at time 0. Wall contact (grey bar) begins between 2-3 seconds and ends between 9-11 seconds, depending on the wall distance and whisker length. Scale bar: 20 passes. (**B**) Peri-event time histograms (PETHs) computed as spike density functions aligned to wall onset time (t=0 s), for the same example units shown in A. Error shading: SEM. (**C**) Z-scored tuning curves (mean ± 95% CI) to wall distance for same example units shown in A, B. (**D**) Heatmap of normalized tuning curves for wall-activated, distance-tuned units (n=472 from 36 mice), sorted by preferred wall distance from close (top) to far (bottom). Tuning curves were linearly interpolated between presented distances; unpresented distances are shown as white space. (**E**) Histogram of preferred wall distances, measured as the location of the tuning curve peak, from units in D. (**F**) Fraction of recorded single units (n=1141) in PrV defined as either untuned (gray), suppressed only (purple), map (yellow, prefer ≥ 8 mm), proximity (pink, prefer < 8 mm), or ambiguous (orange), which could not be statistically defined as either proximity or map (See Supplemental Figure S3).

We identified 656/1,141 (57.5%) units that were tuned to wall distance. Among these distance-tuned units, 472/656 (72.0%)) were significantly activated by at least one distance relative to baseline, 213/656 (32.5%) were significantly suppressed by at least one distance relative to baseline, and 29/656 (4.4%) met both criteria (activated at some distances and suppressed at others). Interestingly, “wall-activated”, “distance-tuned” units exhibited lower baseline firing rates than untuned units (5.46 ± 0.670 vs. 6.90 ± 0.682 sp/s; two-tailed Kolmogorov-Smirnov (K-S) test, *p* = 1.91 × 10^⁻10^), while “wall-suppressed”, “distance-tuned” units exhibited baseline firing rates comparable to untuned units (7.31 ± 1.124 vs. 6.90 ± 0.682 sp/s; two-tailed K-S test, *p* = 0.27; Supplemental Figure S4A). During wall contact, “wall-activated”, “distance-tuned” units showed significant increase in firing relative to baseline (16.44 ± 1.640 vs. 6.36 ± 0.704 sp/s; one-tailed K-S test, *p* = 9.26 × 10⁻^28^; Supplemental Figure S4B, left), “wall-suppressed”, “distance-tuned” units showed significant decrease in firing relative to baseline (4.47 ± 1.000 vs. 6.19 ± 1.076 sp/s; one-tailed K-S test, *p* = 3.06 × 10⁻^4^; Supplemental Figure S4B, right), whereas untuned units showed no difference (6.65 ± 0.687 vs. 6.90 ± 0.682 sp/s; two-tailed K-S test, *p =* 0.74; Supplemental Figure S4C), validating our classification criteria.

To visualize tuning at the population-level, we concatenated normalized tuning curves for wall-activated, distance-tuned units (Figure 2D) and wall-suppressed, distance-tuned units (Supplemental Figure S4D) into heatmaps, sorted by each unit’s preferred wall distance. While many units exhibited tuning to the closest wall position (7 mm), others showed tuning to intermediate or far distances (Figure 2E, Supplemental Figure S4F). To further characterize wall-activated distance-tuned units, we compared mean firing rates at each non-closest distance (≥ 9 mm) to the closest distance (7 mm; one-tailed Wilcoxon signed-rank tests, see STAR Methods). Units with peak firing at the closest distance were classified as “proximity” units (Supplemental Figure S4G, left), while units with significantly higher firing at any non-closest distance were categorized as “map” units (Supplemental Figure S4G, right). This analysis identified 266 proximity units (23.3%) and 118 map units (10.3%) out of the 1,141 total recorded units (Figure 2F). An additional 88 units (7.7%) were tuned but did not meet criteria for either map or proximity and were labeled “ambiguous” (Supplemental Figure S4G, middle) and excluded from further analysis. Suppressed-only units comprised a small portion of the population (119/1141; 10.4%, Figure 2F).

We note that classification as “proximity” versus “map” is based on relative peak firing rates within the tested range (7-23 mm). It remains possible that some proximity units would exhibit decreased firing at even shorter distances, thus revealing map-like tuning beyond the tested range. Therefore, our categorization serves as an operational definition to capture distinct response properties within the presented distance range, rather than strict functional classes.

To assess the reliability of proximity and map codes across repetitions, we performed 50% cross-validation by sorting units using half the wall passes and applying the sorting order to the remaining half. The resulting distance-tuning heatmaps showed little difference in preferred distance between cross-validated partitions (Supplemental Figure S4E). Together, these results reveal that PrV robustly encodes peri-head distance using both a proximity code—characterized by increasing firing at close distances—and a distributed map code. Though the map coding is more prominent in wS1 (Sofroniew & Vlasov et al., 2015), its presence in PrV points to an underappreciated role for the whisker brainstem in active sensory processing, beyond simple relay.

### The map code in PrV allows more accurate readout of peri-head distance

We next asked which encoding scheme, proximity or map, allows more accurate readout of peri-head distance from a given neural population. To address this, we trained maximum correlation coefficient decoders (see STAR Methods) on either the proximity or map subpopulation activity, with untuned and shuffled subpopulations used as controls.

As expected, decoding accuracy during the wall-contact period was higher in both map and proximity populations compared to controls. Notably, the map population (n = 118 units) outperformed the proximity population (subsampled n = 118 of 266 units) in overall distance decoding accuracy over the entire wall passing period averaged across all wall distances (Figure 3A). Untuned units still performed slightly above chance due to residual distance tuning that fell below our stringent statistical threshold for wall-tuning classification.

**Figure 3.**
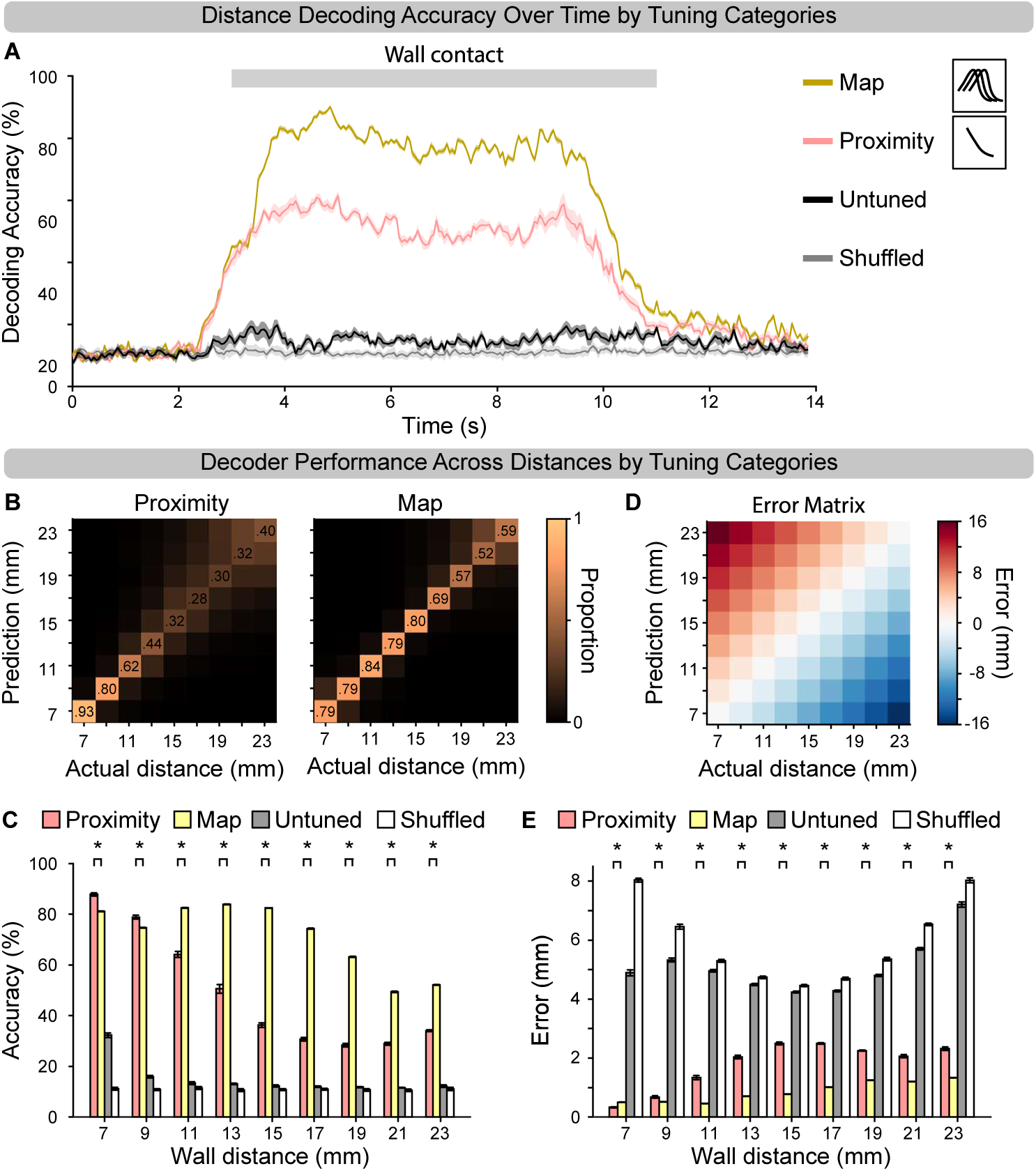
The map code enables more accurate decoding of peri-head distance. (**A**) Decoding accuracy over time averaged across all distances for population decoder models trained on subsamples of 118 units from either proximity (pink, n=266), map (yellow, n=118), untuned (black, n=550), or trial-shuffled control (gray, n=384) subpopulations. Shaded area: 95% confidence interval. Wall contact occurs around 3-11s (gray bar) after the start of wall movement at t=0 s. (**B**) Average confusion matrices showing proportion of wall distance decoded for each actual distance from proximity (left) and map (right) populations. (**C**) Bar plot showing mean ± SEM population decoding accuracy for proximity (pink), map (yellow), untuned (gray), and shuffled control (white); *, *p*<0.05, Wilcoxon rank-sum test with BHFDR correction. (**D**) Matrix quantifying error, in mm, for each coordinate in the confusion matrix. (**E**) Bar plot showing mean decoding error ± SEM in mm for decoders trained on each population (same colors as D); *, *p*<0.05, Wilcoxon rank-sum test with BHFDR correction. Error was computed by multiplying the confusion matrix with the absolute value of the error matrix and summing across rows.

Confusion matrices revealed that map units were particularly more accurate at decoding intermediate and far distances (Figure 3B, right), whereas proximity units showed stronger decoding performance only at the closest distance (Figure 3B, left). To quantify this, we compared classification accuracy across nine wall distances for each population (Figure 3C; Table S2). At 7 and 9 mm, proximity units were significantly more accurate than map units. In contrast, map units were significantly more accurate at decoding more distal wall positions (11-23 mm). To assess decoding precision in millimeters, we computed a distance-classification error (in mm) by multiplying the confusion matrix with a distance error matrix (Figure 3D; Table S3). While classification error increased with distances, both populations showed submillimeter error in decoding the closest distances (7 and 9 mm). However, only the map population showed submillimeter error in decoding intermediate distances (from 11 to 15 mm, Figure 3E). These results are consistent with the performance of single-neuron decoders trained on map and proximity units and averaged across those populations (see STAR Methods, Supplemental Figure S5; Table S4; Table S5), although their overall decoding accuracy is lower than population decoders.

Together, these results suggest that proximity and map codes provide complementary encoding schemes to read out peri-head distance: the proximity code supports highly accurate decoding at close distances, while the map code enables finer discrimination across intermediate and far distances. Thus, optimal distance readout by downstream circuits would integrate information across both proximity and map subpopulations.

### Multi-whisker integration contributes to the generation of the PrV map code

In the wall-passing paradigm, closer wall distances induced greater whisker bending, driving stronger activation of whisker TG afferents. TG therefore transmits to PrV a monotonic, proximity-based input, with firing rates increasing as stimulus distance to the head decreases. The presence of PrV neurons tuned to specific wall distances (i.e., map cells) suggests a transformation of this peripheral proximity code into a representation of peri-head distance. Because the passing wall stimulates multiple whiskers of different lengths, we hypothesized that PrV could generate the map code by integrating multi-whisker inputs. To test this hypothesis, we recorded PrV single-unit activity during the wall-passing paradigm under two conditions within the same recording session: first, with the mouse’s full whisker array intact, and second, after acutely cutting all but a single whisker, thus eliminating multi-whisker integration (see STAR Methods; Figure 4A). Continuous recording of neural activity allowed us to track each neuron’s distance tuning profile before and after the acute whisker cutting manipulation.

**Figure 4.**
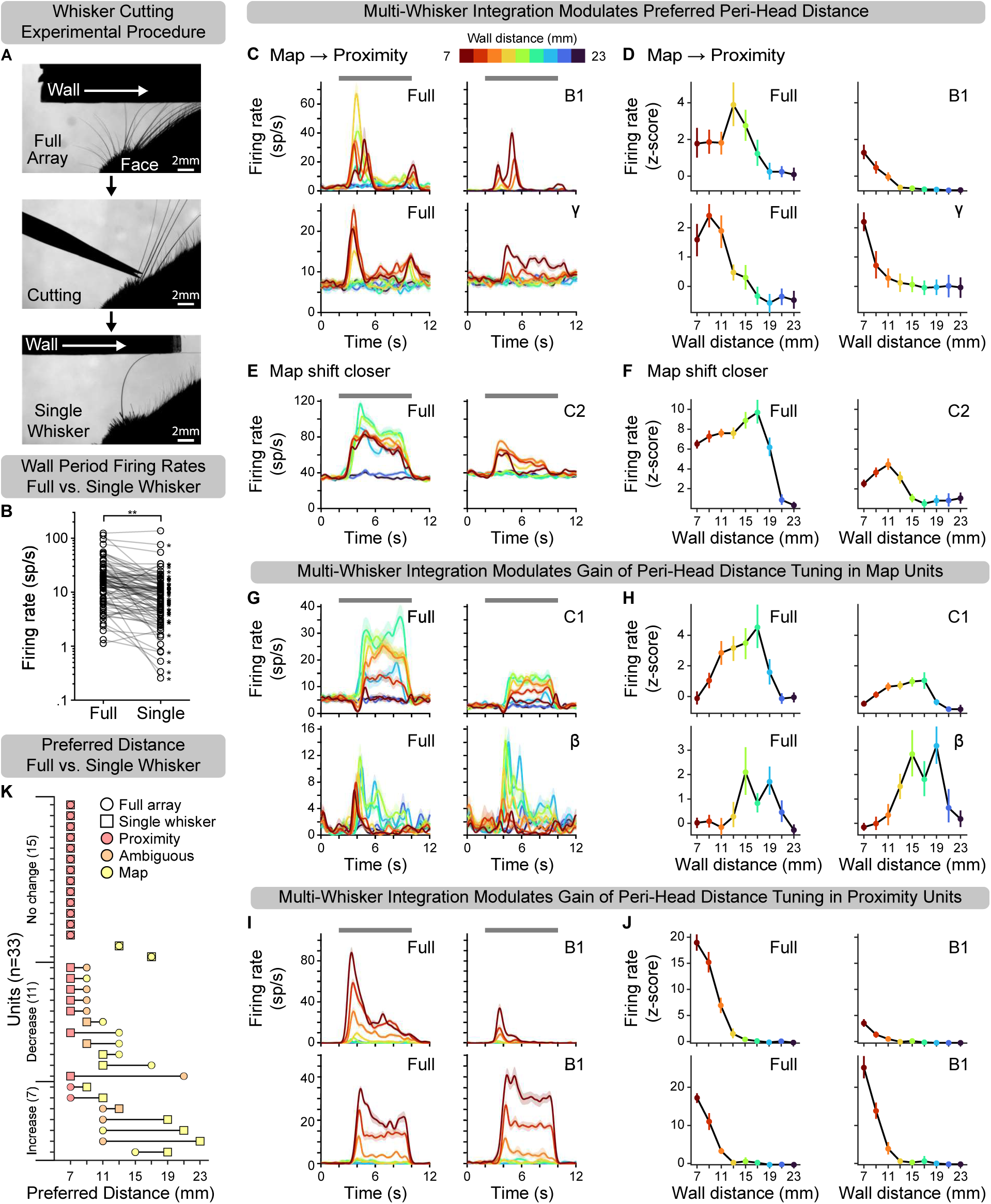
Multi-whisker input supports but is not required for the distance map code. (**A**) Top, Example video frame showing multiple whiskers during wall-passing stimulation (wall moving left to right). Middle, same session during whisker cutting. Bottom, same session after cutting to a single B1 whisker. (**B**) Pairwise comparisons of firing rates (log scale) during wall-passing periods for wall-activated and distance-tuned units (n=86 units from 18 mice) in the full-whisker versus single-whisker condition. Lines connect the same unit across conditions. **, Population mean firing rates: 11.58 ± 2.986 sp/s (full array) vs. 8.18 ± 2.602 sp/s (single whisker), *p* = 4.43 × 10^-8^); *, individual neuron, *p* < 0.05; Wilcoxon signed-rank tests. (**C**,**D**) Two example map units that converted to proximity units after cutting down to a single whisker (B1, top; γ, bottom): (C) Spike density functions (SDFs); (D) corresponding tuning curves. (**E**,**F**) Example map unit that retained map-like tuning but shifted its preferred wall distance closer after cutting down to a single C2 whisker. (**G**,**H**) Two example map units showing gain modulation after cutting down to a single whisker (C1, top; β, bottom): Top, decreased activity; Bottom, increased activity (disinhibition). (**I**,**J**) Two proximity units showing similar gain modulation effects: Top, decreased activity; Bottom, increased activity (disinhibition). (**K**) Pairwise comparison of preferred wall distances for wall-activated units (n=33 units from 13 mice) that remained distance-tuned after cutting down to a single whisker. Lines connect the same unit across conditions. Open circles: map units; closed circles: proximity units. Color indicates tuning category: proximity (red), ambiguous (orange), map (yellow).

Across the recorded population (n = 195 units from 18 mice), acute whisker cutting did not significantly alter baseline firing rates (measured before wall-movement onset for each wall pass; Wilcoxon signed-rank test, full array: 7.09 ± 1.034 vs. single whisker: 7.10 ± 1.038 sp/s, *p* = 0.70). In contrast, firing rates for wall-activated, distance-tuned units (n = 86) during wall-passing periods were significantly reduced following cutting down to a single whisker (full array: 11.58 ± 1.338 vs. single whisker: 8.18 ± 2.602 sp/s, *p* = 4.43 × 10^−8^, Figure 4B). Of the 86 wall-activated and distance-tuned units, 66 showed significant changes in firing rates during wall-passing periods between cutting conditions (two-sided Wilcoxon signed-rank test, *p* < 0.05). Among these, 52 units showed a significant decrease in firing rates, consistent with loss of excitatory input from whiskers within their receptive fields, while 14 units displayed increased firing rates, likely reflecting disinhibition due to removal of surround-whisker inputs (Furuta et al., 2008). Note that since we could not identify the receptive field of each recorded PrV unit on the high channel count probe, and that distance tuning was computed after the recording session, in many cases the principal whisker driving a given neuron of interest was inevitably cut, leading to a near-complete loss of response.

Of 21 units classified as map cells before whisker cutting, 10 of these remained tuned to wall distance after cutting down to a single whisker. After cutting, we observed shifts in preferred wall distance: two map cells shifted their preferred distance from an intermediate to the closest wall distance (Figure 4C,D; population summary in Figure 4K). The whisker cutting manipulation effectively converted these map cells to proximity cells, indicating that in these cases multi-whisker integration was essential for generating the non-monotonic map code for peri-head distance. On the other hand, four map cells shifted their preferred distance closer but did not fully convert to proximity cells (e.g., Figure 4E,F). Two map cells retained their preferred distance after whisker cutting but showed changes in response amplitude (Figure 4G,H). These results showed that map cells could be generated with inputs from just a single whisker although multi-whisker inputs still modulate the gain or the preferred distance tuning in map cells. Together, these findings demonstrated that multi-whisker integration contributes to, but is not required for, transforming proximity-based inputs into a non-monotonic map code in PrV.

We also examined how proximity cells were affected by the loss of multi-whisker input. 45 units were classified as proximity cells before whisker cutting, and only 15 of these remained tuned to wall distance after cutting down to a single whisker. In remaining proximity cells, reducing input to a single whisker changed the gain of their responses: nine units showed decreased activity (e.g., Figure 4I,J, Top), while four exhibited increased activity (e.g., Figure 4I,J, Bottom).

### Inhibition shapes the PrV map code

In addition to integrating inputs from different whiskers, the sensory code transformation occurring in PrV may also involve the subtraction of TG inputs derived from the same whisker but with differing distance sensitivity profiles. Prior research indicates that TG touch neurons exhibit heterogeneity in mechanosensory thresholds as well as in their adaptation properties (Abraira & Ginty, 2013; Furuta et al., 2020). This suggests that TG touch neurons innervating the same whisker may exhibit different proximity-tuning profiles. Whether involving single- or multi-whisker integration, a common feature of the hypothesized transformation model (Figure 1B) is the engagement of inhibitory mechanisms. Next, we aimed to directly test this by combining *in vivo* electrophysiology with optogenetic or chemogenetic silencing approaches.

The intrinsic circuitry of PrV is shaped by two major sources of inhibition: local glycinergic and GABAergic interneurons within PrV itself, as well as profuse long-range GABAergic projections from the interpolar part of the spinal trigeminal nucleus (SpVi; Furuta et al., 2008; Bellavance et al., 2010). To assess the role of local inhibitory neurons in encoding of peri-head distance within PrV, we unilaterally injected AAV-DIO-GtACR (Govorunova et al., 2015; Takatoh & Prevosto et al., 2021) into PrV of vGat-Cre mice (N=2), enabling Cre-dependent expression of the inhibitory opsin GtACR in PrV inhibitory neurons (Supplemental Figure S6A). We then performed acute optogenetic silencing of these local inhibitory neurons while recording single-unit activity in PrV during the wall-passing paradigm (Supplemental Figure S6B,D). For each wall distance, the "Laser OFF" and "Laser ON" conditions were randomly interleaved. Each distance was repeated 15-20 times in each condition to allow the comparison of distance tuning profiles with and without local inhibition.

Among the 112 well-isolated units, 18 were putative inhibitory neurons expressing GtACR, as they were strongly inhibited during the “Laser ON” period (modulation index ≤ −0.5; see STAR Methods; Supplemental Figure S6C,E). Of distance-tuned units (71/112), 17 were opto-inhibited, exhibiting a decrease in firing rates during laser stimulation (modulation index < 0; see STAR Methods), while preserving their distance tuning profile (i.e., proximity or map, data not shown). Sixteen units showed disinhibition under the “Laser ON” condition (modulation index > 0; see STAR Methods). Nonetheless, these opto-disinhibited neurons, regardless of whether they were proximity or map cells, retained their preferred distance (e.g., Supplemental Figure S6F,G). These observations suggest that PrV local inhibitory neurons primarily modulate the gain of individual PrV neurons’ firing rates, rather than altering their distance tuning.

We next investigated the role of long-range internuclear inhibitory inputs from SpVi to PrV. Due to the extensive size of SpVi, optogenetic silencing of its inhibitory neurons using a single optical fiber is impractical. Instead, we employed an AAV strategy to achieve sustained silencing by expressing tetanus toxin light chain (TeLC; Takatoh & Prevosto et al., 2021) to permanently block synaptic transmission. AAV-DIO-TeLC-EYFP was unilaterally injected into SpVi of vGat-Cre mice (N=11) to target inhibitory neurons known to send long-range projections to PrV (Figure 5A; Furuta et al., 2008). Post-hoc histology confirmed TeLC-EYFP expression in SpVi (Supplemental Figure S6H), as well as labeled axon terminals and boutons in PrV originating from SpVi inhibitory neurons (Figure 5B). Note that the large size of SpVi limited viral access to all PrV-projecting neurons, thereby constraining the extent of silencing.

**Figure 5.**
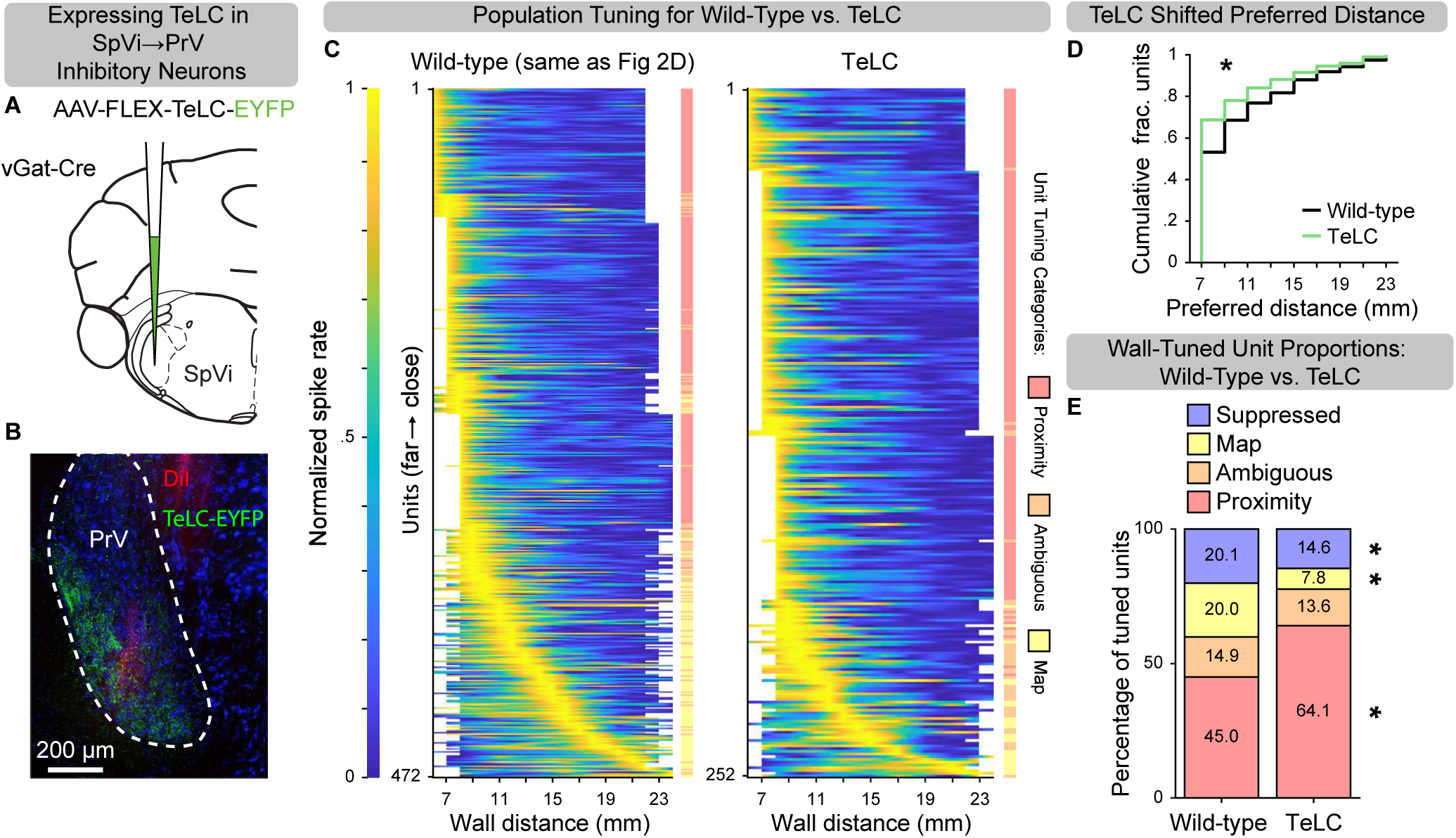
Silencing intersubnuclear inhibitory projections from SpVi diminishes the PrV map code. (**A**) Viral-genetic strategy to express TeLC in inhibitory cells in caudal SpVi, which send long-range projections to PrV. (**B**) Example coronal section at 4.6 mm posterior to Bregma, showing axons of TeLC-labeled inhibitory cells (TeLC-EYFP, green) in PrV and a silicon probe recording track labeled with Dil (red). Cell bodies were stained with Nissl (blue). (**C**) Heatmaps of normalized tuning curves for wall-activated, distance-tuned units in PrV. Left, wild-type mice (same data as Fig. 2D, reproduced here for side-by-side comparison). Right, a separate cohort of vGat-Cre mice with TeLC-mediated silencing of PrV-projecting inhibitory neurons (n=252 units from 11 mice). Units are ordered by preferred wall distance, from near (top) to far (bottom). The color bar to the right of each heatmap indicates unit tuning category. Tuning curves were linearly interpolated between presented wall distances; unpresented distances appear as white space. (**D**) Cumulative distributions of preferred distances for the same units in C, comparing wild-type (black) and SpVi-TeLC (green) populations (*p* = 8.02 × 10^-5^, one-tailed K-S test). (**E**) Comparison of the proportions of distance-tuned units in wild-type (n=1141 from 36 mice) population and in SpVi-TeLC (n=587 from 11 mice) population. TeLC-mediated silencing of PrV-projecting inhibitory neurons in SpVi significantly reduced the prevalence of map units (3.90 vs 10.34%; Z-statistic: 4.64, *p* = 1.80 × 10^-6^, one-tailed Z-test) and suppressed units (7.30 vs 10.43%; Z-statistic: 2.12, *p* = 0.02, one-tailed Z-test), but increased the prevalence of proximity units (32.09 vs 23.31%; Z-statistic: −3.93, *p* = 4.27 × 10^-5^, one-tailed Z-test). The proportions of untuned and ambiguous units were not significantly different between samples from wild-type and TeLC mice. *, significance.

We recorded single-unit activity in PrV 10-15 days after TeLC injection during the wall-passing paradigm and isolated 587 units (Supplemental Figure S3F-I). To assess the impact of suppressing long-range inhibitory inputs from SpVi, we compared wall-distance tuning in SpVi-TeLC mice to that in wild-type mice. Side-by-side comparison of normalized tuning curves of all wall-activated, distance-tuned units revealed a clear reduction in the proportion of map units in SpVi-TeLC mice (Figure 5C, Left: data previously presented in Figure 2, Right: TeLC experiment), shifting the population to prefer closer wall distances (Figure 5D). Tuning profile classification confirmed a significant shift in the distribution of tuning types (Figure 5E). Most importantly, the proportion of map units was reduced by more than half in SpVi-TeLC mice compared to wild-type mice (3.9% vs 10.34%; Z-statistic: 4.64, *p* = 1.8×10^−6^), while the proportion of proximity cells increased (32.09% vs 23.31%; Z-statistic: −3.93, *p* = 4.27×10^−5^). Our silencing manipulation did not fully abolish map cells, possibly due to incomplete silencing of all long-range inhibitory intersubnuclear projections. The proportion of suppressed units decreased modestly (7.3% vs 10.43%; Z-statistic: 2.12, *p* = 0.02), whereas the proportions of untuned (49.9% vs 48.2%; Z-statistic: −0.68, *p* = 0.50) and ambiguous units (6.79% vs 7.71%; Z-statistic: 0.69, *p* = 0.49) remained unchanged.

During both baseline and wall-passing periods, untuned units in SpVi-TeLC mice exhibited significantly higher firing rates than those in the wild-type population, consistent with disinhibition following loss of inhibitory inputs from SpVi (baseline: 9.34 ± 1.330 vs 6.90 ± 0.682 sp/s, *p* = 7.63 × 10^-5^; wall passes: 9.04 ± 1.390 vs. 6.65 ± 0.687 sp/s, *p* =1.06 × 10^-4^; one-tailed K-S tests; Supplemental Figure S6I). In contrast, distance-tuned units in SpVi-TeLC mice showed comparable baseline activity (in the absence of wall contact) to those in wild-type mice (5.60 ± 0.845 vs. 5.46 ± 0.670 sp/s; one-tailed K-S test, *p* = 0.14), but significantly elevated firing rates during wall contact (20.25 ± 2.263 vs. 16.44 ± 1.640 sp/s; one-tailed K-S test, *p* = 1.64 × 10^-3^ Supplemental Figure S6J). These findings suggest that TeLC-mediated silencing of SpVi inputs altered the tuning preferences of PrV neurons (i.e., shifting them toward closer wall distances) without affecting their spontaneous activity level. Thus, long-range inhibition from SpVi plays a key role in transforming feedforward proximity-based inputs from TG into a map-like peri-head distance representation in PrV. More broadly, our findings extend the role of SpVi-PrV inhibitory projections beyond simple sensory gating (Furuta et al., 2008; Mohar et al., 2013), implicating them in a sensory code transformation.

Finally, to demonstrate how this SpVi-PrV inhibitory circuit enables the proximity-to-map transformation, we implemented a simple model constrained by known anatomical motifs (Figure 6A; see STAR Methods). In our model, TG afferents send proximity-based inputs to neurons in PrV via two distinct pathways: a direct excitatory (lemniscal) pathway and an indirect inhibitory pathway mediated by SpVi inhibitory neurons. TG proximity-based inputs were modeled as sigmoidal functions with thresholds skewed toward proximal distances (Figure 6B), and PrV neurons received sparse, random combinations of excitatory inputs from TG neurons and inhibitory inputs from SpVi (see STAR Methods). The model transformed monotonic inputs into a heterogeneous population of PrV tuning curves, including both proximity- and map-like responses to distance (Figure 6C). Across the modeled population (n = 1,225), PrV neurons exhibited a wide range of preferred distances, reproducing qualitative features of the empirical tuning curves (Figure 6D-F, left). Reducing inhibitory input per PrV neuron by 50% led to noticeable loss of map units and expansion of proximity units (Figure 6D-F, right). These simulations show that simple feedforward motifs can transform monotonic sensory input into a non-monotonic map-like code, enabling extraction of an abstract representation of peri-head distance, and that inhibition is necessary for generating intermediate and far distance tuning in PrV. Together, these results highlight complementary roles for local and long-range inhibition in the brainstem trigeminal complex: local inhibitory circuits primarily regulate sensory gain, whereas long-range SpVi inputs reshape tuning to generate the map code for peri-head distance.

**Figure 6.**
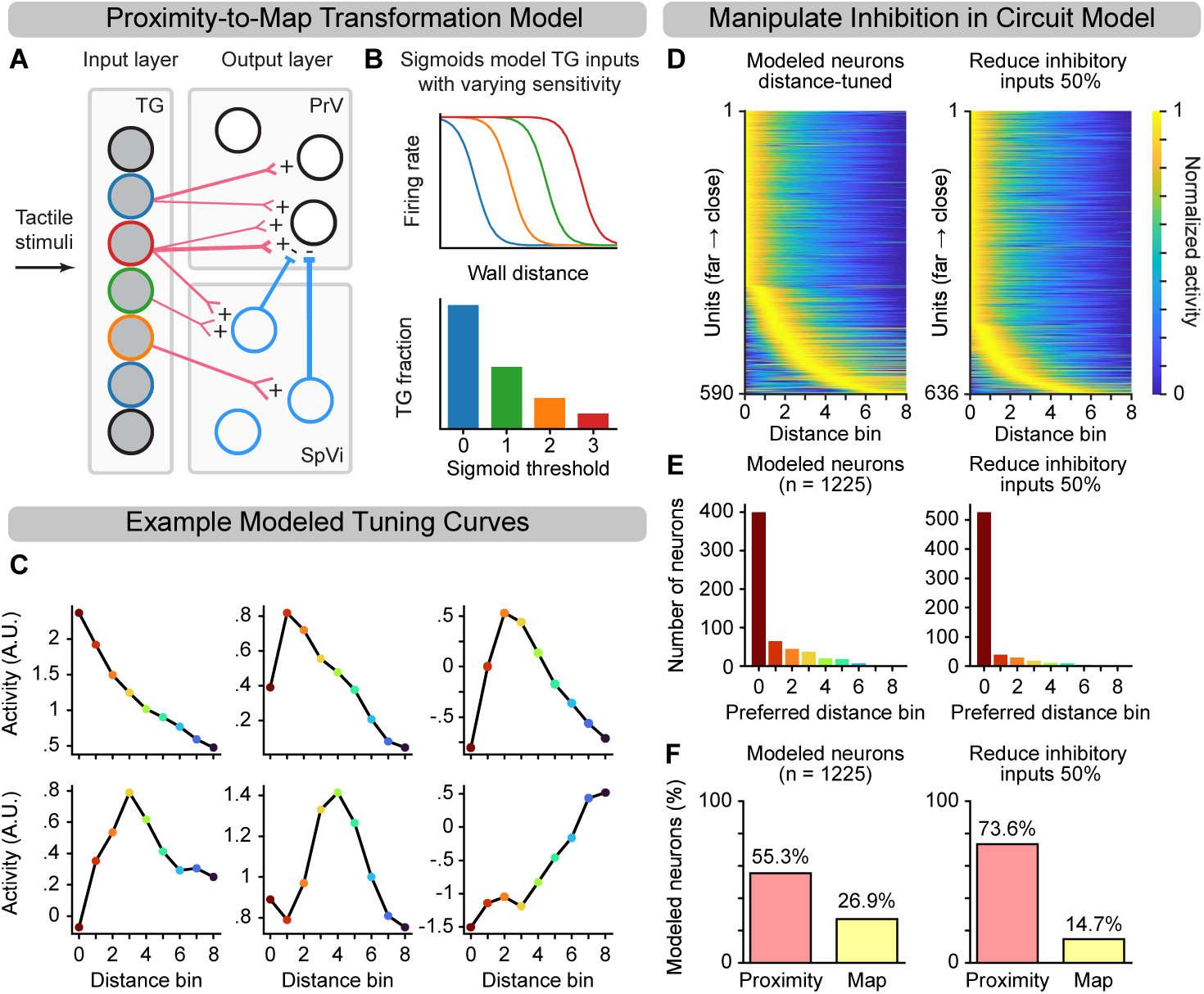
Simulation of brainstem circuitry to generate distance-map code. (**A**) Schematic diagram of the modeled neural circuit. The input layer consists of a pool of TG neurons (gray-filled circles), whose tuning to wall distance was modeled as sigmoidal functions with varying thresholds (indicated by the outline color of each circle). These TG neurons project to PrV neurons (black circles, output layer) via two distinct pathways: a direct excitatory (lemniscal) pathway (red) and an indirect inhibitory pathway (blue). The inhibitory pathway is mediated by a population of inhibitory neurons located in SpVi (blue circles), which project to PrV neurons. (**B**) Top, TG tuning curves (color-coded to match TG neurons outlined in A) are modeled as sigmoidal functions with thresholds spanning the full range of wall distances. Bottom, The sigmoid threshold distribution is exponentially skewed to emphasize the encoding of proximal wall distances. (**C**) Example tuning curves of modeled PrV neurons. Y-axis: baseline-subtracted activity predicted by the model (A.U.). Wall distances are normalized between 0 and 1, and nine uniformly spaced distance bins (labeled 0-8) are used for analysis and visualization. (**D**) Left, Heatmap of normalized tuning curves for distance-tuned units (n=590) in the modeled PrV population, sorted by preferred distance bin from close (top) to far (bottom). Right, Same analysis after reducing inhibitory inputs to each PrV neuron by 50%, n=636 distance-tuned units. (**E**) Left, Histogram of preferred distance bins, defined as the peak (max) of the tuning curve, for modeled PrV neurons. Right, Same analysis after reducing inhibitory inputs to each PrV neuron by 50%. (**F**) Left, Proportion of modeled PrV neurons classified as proximity (pink, prefer distance bin 0) or map (yellow, prefer > distance bin 0). Right, Same classification following 50% reduction of inhibitory inputs per modeled PrV neuron.

## DISCUSSION

Here, we revealed a previously unrecognized sensory code transformation in the whisker brainstem: from proximity-based whisker sensory inputs to map-like peri-head distance representation. Using *in vivo* electrophysiological recordings from awake, behaving mice in a naturalistic wall-passing paradigm, we discovered two encoding schemes for distance in PrV: a proximity code in which neuronal firing rates increase monotonically as the distance becomes closer, and a map code in which neurons exhibit peak firing at specific distances. Both encoding schemes are robust and stable, invariant to the multiplexed and highly variable whisker inputs across trials. Mechanistically, we provide evidence that both multi-whisker integration and long-range inhibitory inputs from SpVi play critical roles in generating the non-monotonic map code. While most prior research on whisker-based object localization has examined neural activities of TG afferents (Ahissar & Knutsen, 2008) and cortical areas (Barnéoud et al., 1991; O’Connor et al., 2010; Sofroniew & Vlasov et al., 2015) in awake, behaving rodents, the brainstem trigeminal complex has remained largely unexplored. To our knowledge, only two prior studies have reported *in vivo* extracellular single-unit activity in PrV (Moore et al., 2015; Chakrabarti & Schwarz, 2018). Our findings thus fill a critical gap in understanding early stages of tactile processing, revealing an anatomically-defined brainstem circuit that integrates and transforms peripheral signals to construct representations of peri-head space. More broadly, this work highlights the role of PrV as a processing center that converts afferent inputs into meaningful representations for downstream computation.

### Functional roles of proximity and map codes in PrV

Our discovery of PrV map cells challenges a long-held assumption of PrV as a simple relay of trigeminal input. The map code, in which neurons fire maximally at a preferred distance and decrease their response as the stimulus deviates from that value, mirrors a common encoding motif observed across sensory systems (Porter & Groh, 2006). Such “peaked” tuning could be computationally advantageous, as it allows distance information to be extracted from a relatively small subset of neurons (Pouget et al., 2000; Jazayeri & Movshon, 2006), resulting in a sparse, metabolically efficient representation capable of attaining high precision (Laughlin, 2001; Olshausen & Field, 2004). The presence of a map code in the brainstem allows feature extraction from multiplexed tactile signals during active sensation (e.g., contact forces, stick-slips, vibrations), reducing computational burden for downstream structures and enabling rapid extraction of behaviorally relevant features.

Functionally, the map and proximity codes could be specialized for different operating regimes of PPS. Our decoder analysis showed that map cells were significantly better at decoding intermediate and far distances, suggesting that they are optimized for fine-grained spatial localization over a broader distance range within PPS. This fine-grained distance representation is likely critical for tasks in which rodents use their whiskers to judge whether gaps are crossable (Azarfar et al., 2018), detect apertures just large enough to pass through (Harris et al., 1999; Grant et al., 2018), follow walls at a consistent distance (Sofroniew et al., 2014; Sofroniew & Vlasov et al., 2015), or avoid obstacles during locomotion (Sofroniew et al., 2014; Warren et al., 2021). The presence of map cells in the brainstem may significantly enhance the speed and precision of these sensorimotor computations. Consistent with this idea, PrV projects directly to the superior colliculus (Bosman et al., 2011), providing a subcortical pathway for distance signals from the brainstem to premotor circuits that bypasses the thalamus and cortex. This direct routing could support particularly rapid sensorimotor coordination, enabling swift orienting, approach, or avoidance responses based on peri-head distance signals. Additionally, map cells could be better suited for 3D shape perception, allowing the brain to detect small variations in contact distance across the whisker array to reconstruct an object’s surface geometry. In contrast, proximity cells excel at decoding the closest distance, making them well suited for threshold-like detection of imminent contact in peri-head space and for triggering rapid, reflexive responses, while simultaneously encoding distance-invariant features such as texture, softness, or compliance.

While proximity and map cells may lie on a continuum of distance tuning rather than forming strictly discrete classes, the distinction provides a useful framework for comparing representational strategies along the lemniscal pathway. In TG, radial distance encoding is mechanically driven and predominantly proximity-based (Szwed et al., 2006; Ahissar & Knutsen, 2008; Pammer & O’Connor et al., 2013; Birdwell et al., 2007; Bush et al., 2016). In PrV, we identified the presence of map cells, but proximity cells are overrepresented. In wS1, a more balanced distribution of distance preferences was reported (Sofroniew & Vlasov et al., 2015). This gradient suggests a progressive transformation from low-level mechanical inputs to higher-order, feature-selective representations of peri-head space that are increasingly aligned with the demands of sensorimotor behavior.

### Feedforward inhibition transforms proximity inputs into a map code

The results from SpVi-TeLC experiments and computational modeling support a canonical E-I-E motif—monosynaptic excitation from TG combined with long-range disynaptic inhibition from SpVi—that converts monotonic, proximity-based TG input into non-monotonic, distance-selective responses in PrV map cells. Specifically, inhibitory inputs from SpVi act as “neural comparators”: by subtracting proximity signals with different distance sensitivity profiles, they allow PrV map cells to exhibit peak activity at specific distances. Our findings extend beyond prior work showing that SpVi inhibition sharpens angular tuning in PrV (Bellavance et al., 2010)—a form of gain modulation—by demonstrating that the same projection can fundamentally change how features are encoded, revealing a broader function for disynaptic inhibition in sensory code transformation. More broadly, these findings situate the SpVi-PrV pathway within a growing class of microcircuits involving inhibitory neurons that implement core sensory computations. Across modalities, feedforward inhibition does not simply limit excitation but actively extracts features by implementing subtractive comparison operations to transform stimulus representations. Examples include binaural circuits in the lateral superior olive of the auditory brainstem, where excitatory and inhibitory inputs carrying sound level information from the two ears are compared to encode interaural level differences (Benichoux & Tollin, 2018), inhibitory subnetworks that sculpt orientation and higher-order tuning in visual cortex (Niell & Scanziani, 2021), and lateral inhibition in olfactory circuits that decorrelates odor representations (Geramita et al., 2016). Recent connectomic analyses have revealed that inhibitory neurons form stereotyped, feature-targeted connectivity patterns (Schneider-Mizell et al., 2025), supporting the idea that inhibition is a commonly repeated anatomical substrate poised to perform computations underlying neural code transformations. By showing that this convergent E-I-E motif (Figure 1B; Figure 6A) in the first central processing center of the whisker system can transform monotonic inputs from the periphery into a discrete distance map, our work supports a unifying view of this motif as a computational primitive for constructing higher-order sensory representations.

### Map code generated with and without multi-whisker integration

We initially hypothesized that the PrV map code emerges through integration of inputs from multiple whiskers. Our whisker cutting experiments showed that a subset of PrV map cells retained map-like distance tuning even when only a single whisker remained. These observations indicate that multi-whisker integration supports, but is not required for, the generation of map-like tuning in a subset of PrV neurons. Together with the diversity of mechanosensory thresholds among TG touch neurons (Hao & Delmas, 2010; Furuta et al., 2020), these observations refined our circuit model whereby PrV constructs the map code by subtracting inputs from TG afferents (from same or different whiskers) with different proximity-tuning profiles.

Our study does not rule out peripheral mechanisms that may contribute to the map code. Mechanosensory receptors in the whisker pad skin (Rice et al., 1986; Severson et al., 2019), including low-threshold mechanoreceptors (LTMRs) that encode skin stretch (Abraira & Ginty, 2013; Grigg, 1996) and C-LTMRs in the surrounding hairy skin (Li et al., 2011), could convey additional distance information to PrV. These signals may encode whisker pad tension and its dynamic changes during wall contact, thereby shaping distance representations in ways not captured by our current understanding of TG input. Notably, conventional extracellular recordings in TG predominantly sample large-diameter neurons, making it technically difficult to assess the contribution of C-LTMRs and other small-diameter afferents. Future work will be necessary to disentangle the relative contributions of central and peripheral mechanisms in constructing peri-head distance representations.

## AUTHOR CONTRIBUTIONS

F.W. conceived the study. F.W., K.S.S, V.P., and W.X. designed the experiments. W.X. and K.S.S. performed all experiments. P.M.T. and V.P. provided training and guidance to W.X. on in vivo extracellular recording in the brainstem. K.S.S. and V.P. developed analysis code with input from W.X. H.Z. implemented the brainstem circuitry simulation. K.C. and M.L. advised on data analysis and statistical methods. J.T advised on viral constructs and viral injection surgeries. K.S.S., W.X., and V.P. analyzed the data. S.Z. generated GtACR and TeLC viruses. S.C. managed the mouse colony, performed genotyping, and bred all animals. W.X., K.S.S, and F.W. wrote the manuscript with input from all authors.

## ACKNOWLEDGMENTS

We thank Drs. Jennifer Groh, David Kleinfeld, Lindsey Glickfeld, and John Pearson, as well as members of the Wang Lab, for insightful discussions of this study. We are grateful to Julia Dziubek, Ina Bando, and He Jiang for technical assistance, and to Priyadarshini Dutta and Gabrielle Sevrain for assistance with mouse colony maintenance. This work was supported by grants NS077986, DP1 NS137188, and a subcontract of U19NS137920 (to F.W.).

---

**Figure S1.**
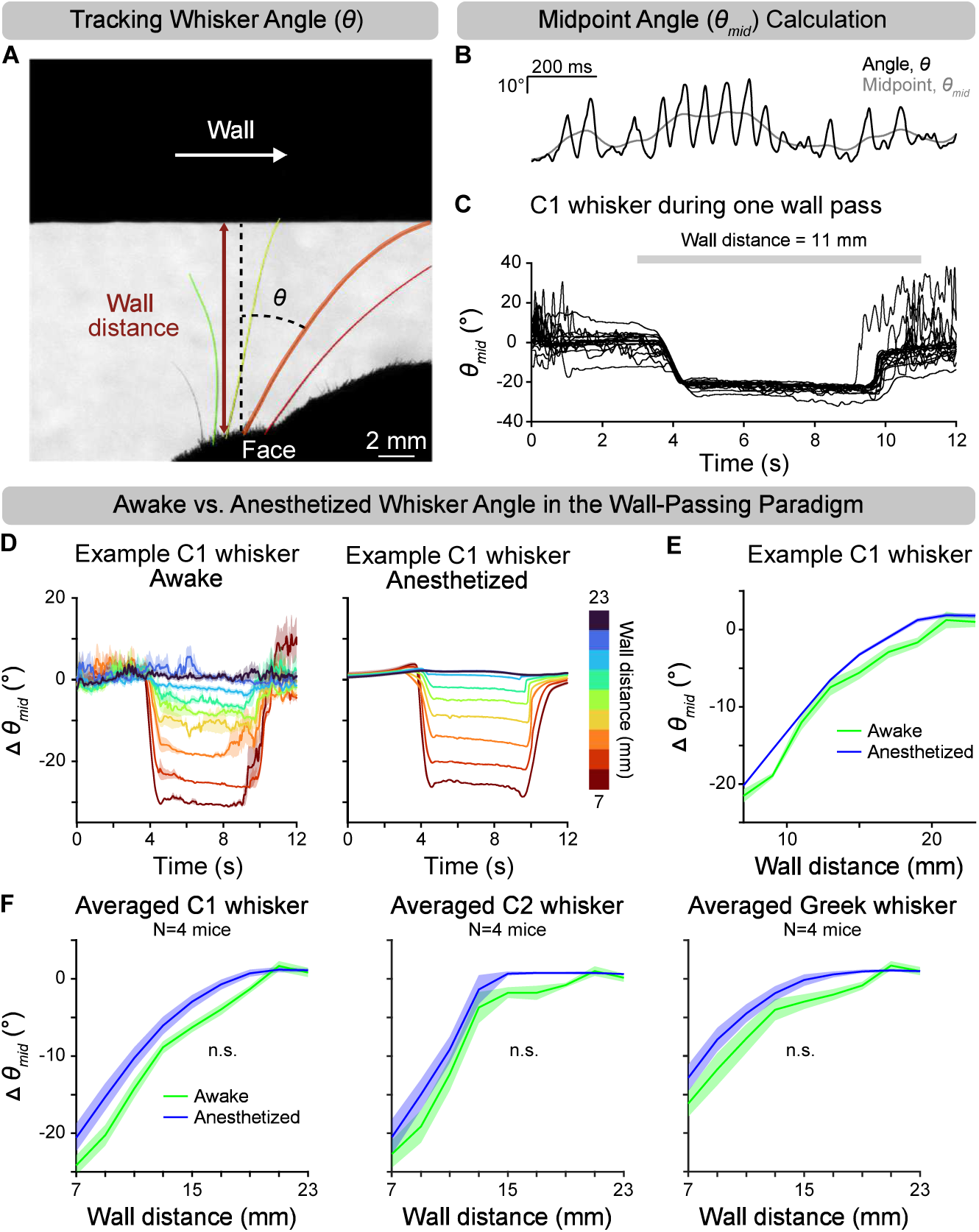
Wall-passing stimulus induced minimal active whisker retraction. (**A**) In a subset of mice (N=4), high-speed video was acquired during the wall-passing paradigm under both awake and anesthetized conditions. Example video frame with tracked whiskers color-coded left-to-right of an awake, behaving mouse: C3 (green), C2 (yellow), C1 (orange), and y (red). Whisker angle (*θ*) was measured relative to the mediolateral axis (dotted line); example shown for C1 (orange). (**B**) Example trace of whisker angle (*θ*, black) and its midpoint (*θ_mid_*, gray) from an awake, behaving mouse; *θ_mid_* is the slowly varying centerline of *θ.* (**C**) Baseline-subtracted midpoint angle for the C1 whisker of 20 repetitions of one wall distance, aligned to wall movement onset (t=0 s) in an awake, behaving mouse. Baseline was defined during the pre-wall period. Gray bar: wall contact period (3-11 s). (**D**) Comparison of whisker midpoint angle between awake and anesthetized conditions in the wall-passing paradigm. Peri-event time histograms (PETHs) show changes in baseline-subtracted midpoint angle (*Δθ_mid_*) for C1, aligned to wall onset (t=0 s) in one mouse. Left: awake, behaving; right: anesthetized. Each trace is the mean of 20 repetitions of each wall distance (color) ±SEM shading. (**E**) Mean change in *Δθ_mid_* versus wall distance for the same C1 whisker shown in D. Green: awake, behaving; blue: anesthetized. (**F**) Same as E, but averaged across mice (N=4). Subpanels show mean *Δθ_mid_* versus wall distance for left: C1, middle: C2, right: Greek whiskers. Lines indicate across-mouse means ±SEM. Green: awake, behaving; blue: anesthetized. Significance reflects mouse-level paired comparisons (awake vs. anesthetized) with Bonferroni correction (*α* = 0.017): C1, *p* = 0.03 (n.s.); C2, *p* = 0.04 (n.s.), Greek: *p* = 0.0199 (n.s.).

**Figure S2.**
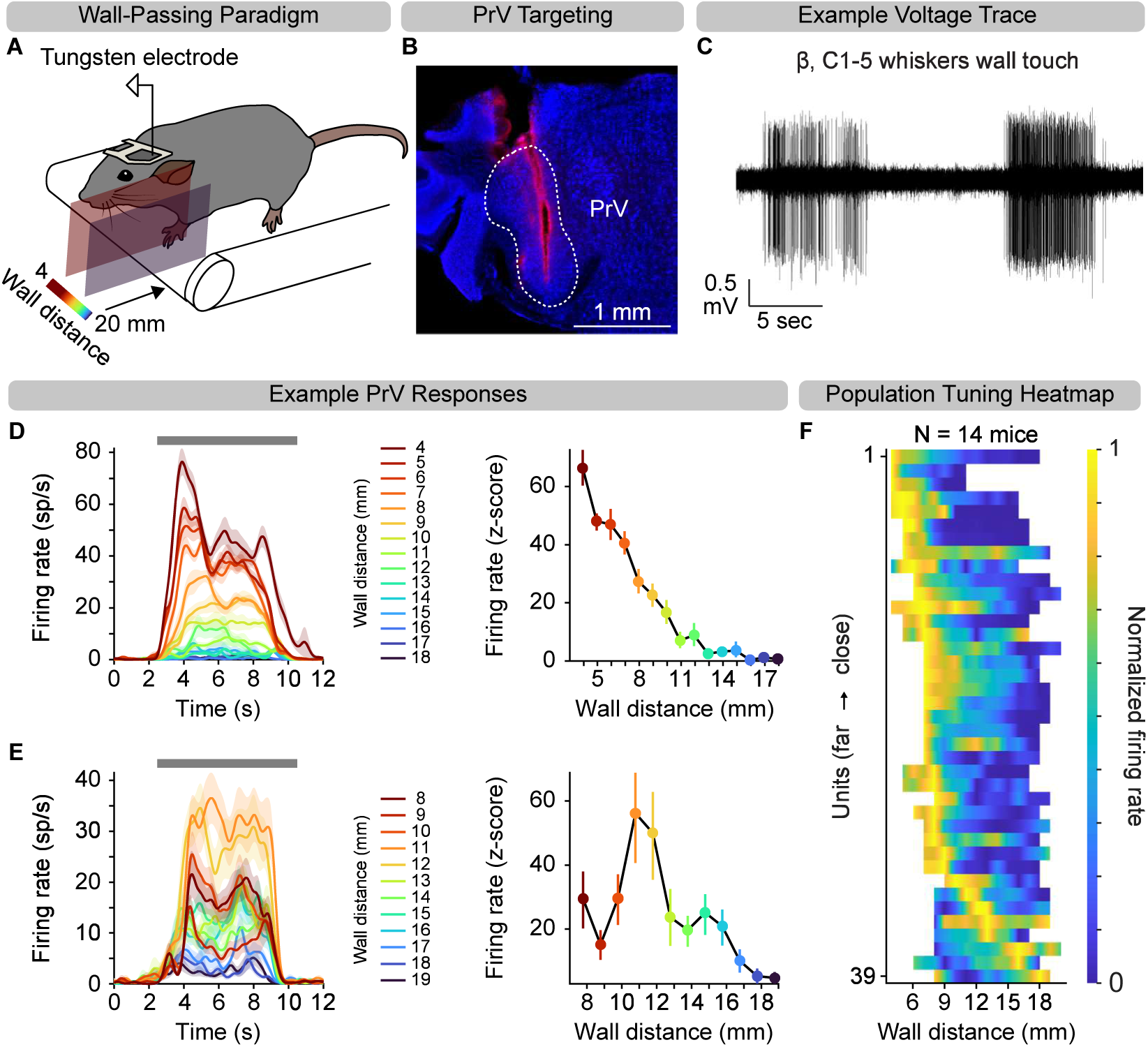
Acute single-electrode recordings from PrV during the wall-passing paradigm. (**A**) Schematic of the experimental setup for acute single-unit recordings in PrV using tungsten electrodes. The wall stimulus moved past the mouse’s face at randomized distances ranging from 4 to 20 mm in 1 mm increments (10-20 repetitions per distance). The mouse has C-row whiskers intact. (**B**) Example coronal brain section at 4.8 mm posterior to Bregma showing one recording track through PrV labeled with Dil (red). Cell bodies were stained with Nissl (blue). (**C**) Example extracellular voltage trace from one recording session, showing two wall pass events. (**D**) Left, PETHs computed as spike density functions aligned to wall onset time (t=0 s), for an example PrV unit preferring the closest distance presented. Error shading: SEM. Right, Z-scored tuning curve (mean ± 95% CI) to wall distance for the same unit. (**E**) Same as D, but for a PrV unit preferring an intermediate distance. (**F**) Heatmap of normalized tuning curves for all distance-tuned units (n=39 from 14 mice), sorted by preferred distance from close (top) to far (bottom). Tuning curves were linearly interpolated between presented distances; unpresented distances are shown as white space.

**Figure S3.**
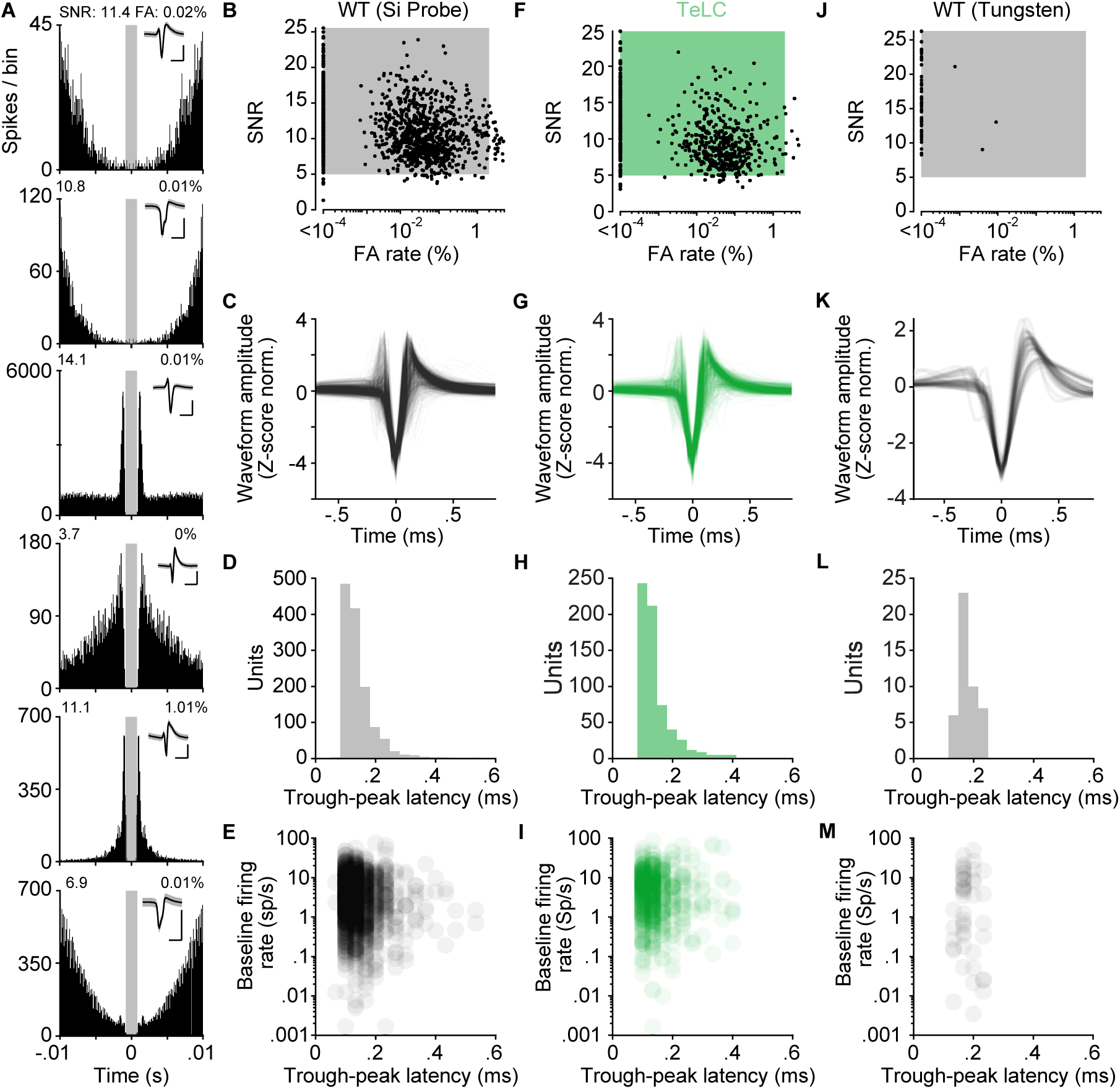
Spike sorting quality metrics. (**A**) Inter-spike-interval (ISI) histograms for the example units in Figure 2A-C. The grey shaded region indicates ± 0.8 ms refractory period. Insets show the corresponding mean spike waveforms; shaded areas represent ± STD. Scale bars: 0.5 ms (horizontal), 0.25 mV (vertical). (**B**) Scatter plot of waveform signal-to-noise ratio (SNR) versus ISI false alarm (FA) rate for wild-type dataset collected with silicon probes (WT-Si Probe; n=1358 total units). The area within the grey square contains acceptable units with SNR > 5 and false alarm rate < 2% (n=1301 units from 36 mice). (**C**) Z-scored waveforms of a subset of acceptable units from the WT-Si Probe dataset (n=100 waveforms shown). (**D**) Histogram of spike widths of all acceptable units from the WT-Si Probe dataset. (**E**) Scatter plot of spike widths, in ms, versus mean firing rate, in spikes per second, for baseline periods (prior to wall pass) of acceptable units. (**F-I**) Same as (B-E), but for tetanus toxin (TeLC) dataset (n=665 total units from 11 mice; n=631 acceptable units in green shaded area). (**J-M**) Same as (B-E), but for tungsten dataset (n=46 total units from 14 mice; n=46 acceptable units in grey shaded area; n=46 waveforms shown in K).

**Figure S4.**
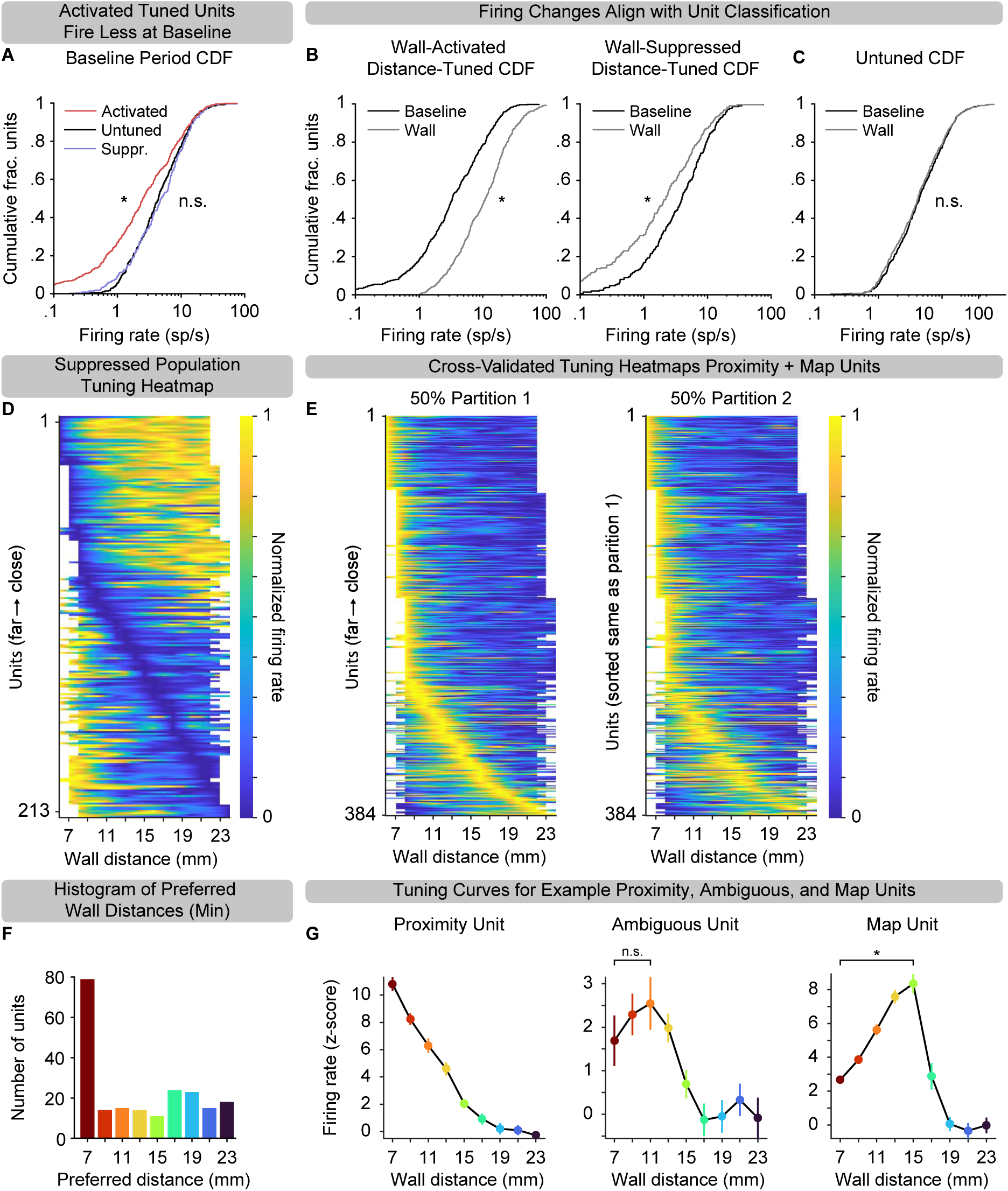
Co-existence of proximity and map codes for encoding peri-head distance. (**A**) Cumulative distribution functions (CDFs; log scale) of baseline firing rates for wall-activated, distance-tuned units (red), wall-suppressed, distance-tuned units (blue), and untuned units (black). Wall-activated units had significantly lower baseline firing rates than untuned units (5.46 ± 0.670 vs. 6.90 ± 0.682 sp/s, mean ± SEM; *p* = 1.91 × 10^-10^, two-tailed K-S test), while wall-suppressed units had firing rates comparable to untuned units (7.31 ± 1.124 vs. 6.90 ± 0.682 sp/s, mean ± SEM; *p* = 0.27, two-tailed K-S test). (**B**) Left, CDFs of firing rates for wall-activated, distance-tuned neurons during wall passes (red) versus baseline periods (black). Firing rates were significantly higher during wall contact at the preferred distance (16.44 ± 1.640 vs. 6.36 ± 0.704 sp/s, mean ± SEM; *p* = 9.26 × 10^-28^, one-tailed K-S test). Right, CDFs of firing rates for wall-suppressed, distance-tuned neurons during wall passes (blue) versus baseline periods (black). Firing rates were significantly lower during wall contact at the preferred distance (4.47 ± 1.000 vs. 6.19 ± 1.076 sp/s, mean ± SEM; *p* = 3.06 × 10^-4^, one-tailed K-S test). (**C**) CDFs of firing rates for untuned units during wall passes (gray) and baseline periods (black). No significant difference was observed (6.65 ± 0.687 vs. 6.90 ± 0.682 sp/s, mean ± SEM; *p* = 0.74, two-tailed KS test). (**D**) Heatmap of normalized tuning curves for suppressed units (n=213), sorted by preferred wall distance from close (top) to far (bottom). Units that were both activated and suppressed appear both in here and in Figure 2D. Tuning curves are linearly interpolated between distances presented and omitted (white space) for distances not presented. (**E**) Cross-validated heatmaps of normalized tuning curves for activated units designated as either “map” (n=118) or “proximity” (n=266), for which trials were randomly partitioned into two equal partitions. Units in left panel were sorted by their preferred wall distances (close-top, far-bottom). Units in the right panel were sorted using the same order as the left panel. (**F**) Histogram of the location of tuning curve troughs from suppressed units in D. (**G**) Z-scored tuning curve (mean ± 95% CI) to wall distance for an example proximity unit (left), ambiguous unit (middle) determined by (i) not significant on the map test (one-tailed Wilcoxon signed-rank test, *p* = 0.09; n.s., not significant), and (ii) not having the highest firing rate at the closest distance, and map unit (right) determined by one-tailed Wilcoxon signed-rank tests comparing firing at each non-closest distance to the closest distance (*p* = 1.27 × 10^-4^; *, *p* < 0.05 with BHFDR correction).

**Figure S5.**
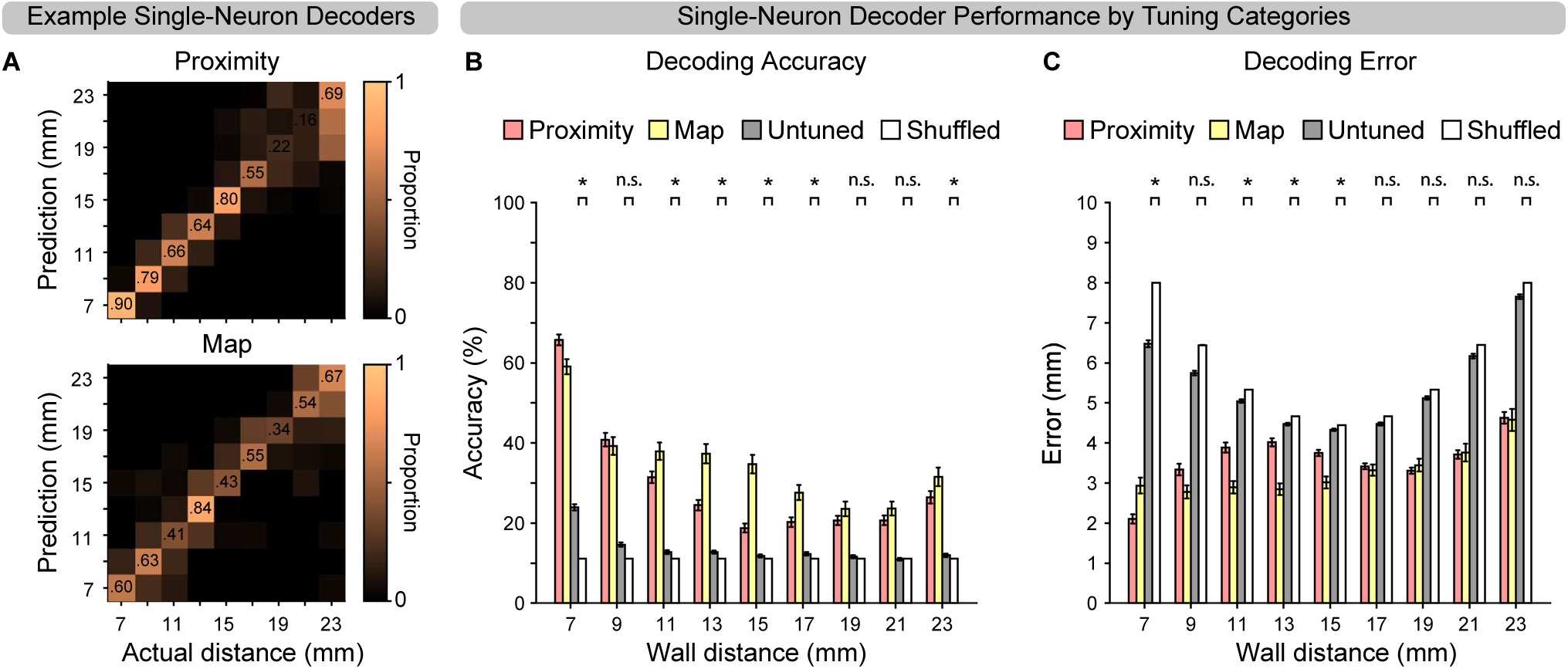
Single-neuron decoding performance. (**A**) Confusion matrices showing proportion of predicted wall distances for each actual distance from two example single neurons. Top, Example proximity unit (Unit 1 in Fig. 2A-C). Bottom, Example map unit (Unit 3 in Fig. 2A-C). (**B**) Mean decoding accuracy (± SEM) of single-neuron decoders grouped by tuning category: proximity (pink, n=266), map (yellow, n=118), untuned (gray, n=550), and trial-shuffled control (white, n=1141). *, *p* < 0.05; n.s., not significant (Wilcoxon rank-sum test with BHFDR correction). (**C**) Mean decoding error (± SEM) in mm by tuning category (same color scheme as B). Error was computed as the elementwise product of the confusion matrix and absolute error matrix (Fig. 3D), summed across rows. *, *p* < 0.05; n.s., not significant (Wilcoxon rank-sum test with BHFDR correction).

**Figure S6.**
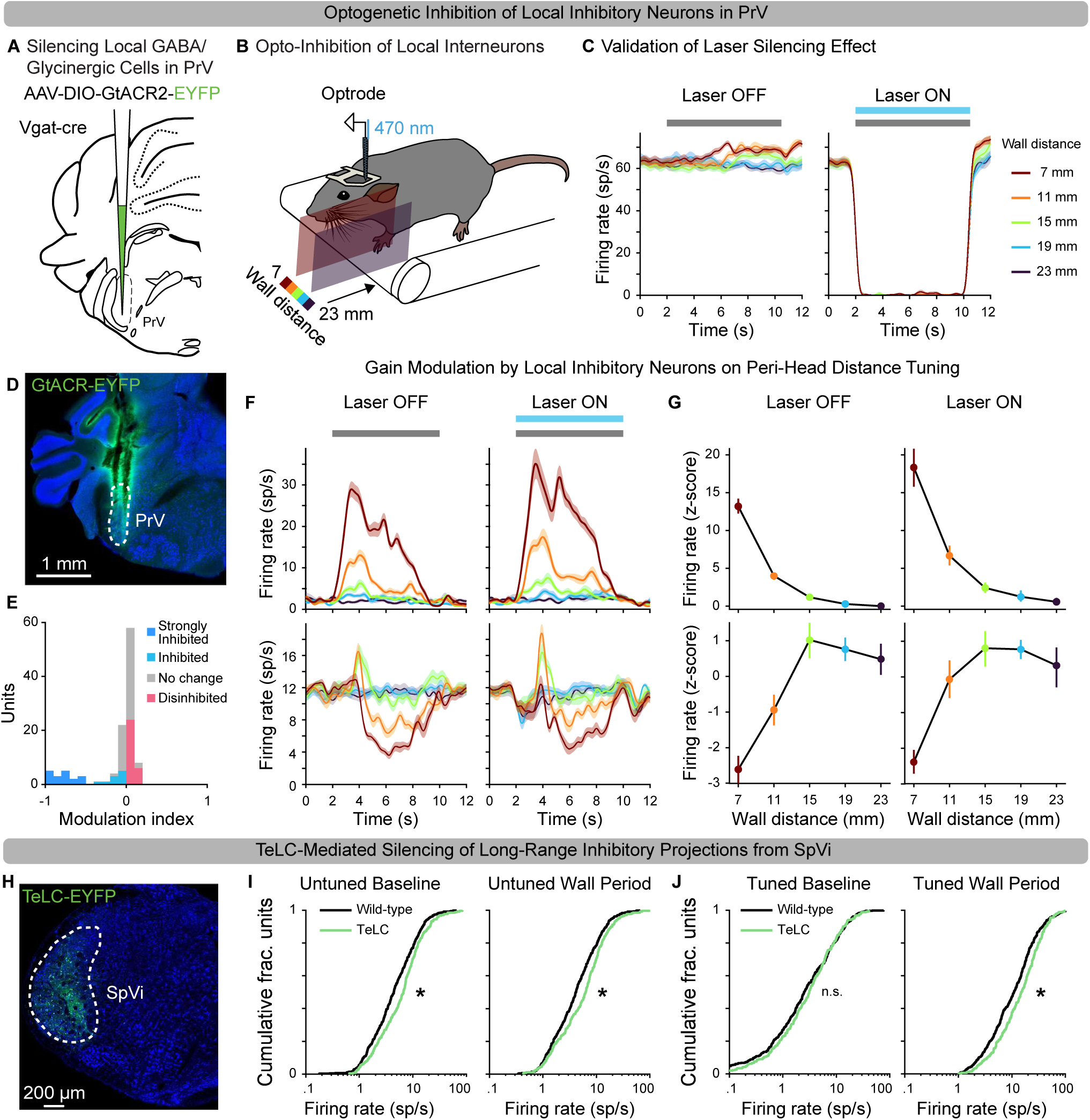
Inhibitory circuit mechanisms contribute to the peri-head distance map code. (**A**) Schematic for expression of the inhibitory opsin, GtACR, in local inhibitory cells in PrV. (**B**) Schematic of acute silicon probe recording paired with optogenetic inhibition of local inhibitory interneurons in PrV. Continuous light stimulation (470 nm, 2 mW) was applied during the entire wall pass with 5s inter-trial interval and randomized laser on/off trials. (**C**) SDFs (in spikes/second) aligned to wall onset time of an example putative inhibitory neuron displaying laser silencing effect during wall passes for “Laser OFF” (left) and “Laser ON” (right) trials. Error shading: SEM. (**D**) Example coronal section at 5.1 mm posterior to Bregma, showing GtACR-labeled inhibitory cells (green) and two silicon probe tracks through PrV, outlined with white dashed line. Cell bodies were stained with Nissl (blue). (**E**) Distribution of modulation index (ML) for all acceptable units (n=112) in the GtACR dataset. Units were classified as strongly inhibited (MI ≤ −0.5, dark blue), inhibited (−0.5 < Ml < 0, light blue), disinhibited (MI > 0, red) when their bootstrap MI confidence interval excluded zero, or as no change when the confidence intervals included zero (gray). (**F**) Top, SDFs of an example proximity unit displaying laser-induced disinhibition in Laser ON (right) condition. an example map unit. (**G**) Z-scored tuning ces for the same units in F. Error bars: 95% Cl. (**H**) Example coronal sectn at 7.4 mm posterior to Bregma, showing the TeLC-labeled inhibitory cells (TeLC-EYFP, green) in SpVi, outlined with white dashed line. Cell bodies were stained with Nissl (blue). (**I**) Left, CDFs of baseline firi rates for untuned units in SpVi-TeLC mice (green) and wild-type mice (black). Firing rates were significantly higher in SpVi-TeLC mice (9.34 ± 1.330 vs. 6.90 ± 0.682 sp/s; *p* = 7.63 x 10^-5^, one-tailed K-S test). Right, CDF the same populations during wall passes. Untuned units in SpVi-Te elevated firing compared to wild-type mice (9.04 ± 1.390 vs. 6.65 ± 0.687 sp/s; one-tailed K-S test, *p* = 1.06 x 10^-4^). (**J**) Left, CDFs of baseline firing rates for wall-activated, distance-tuned units in SpVi-TeLC mice (green) and wild-type mice (black). No significant difference was observed (5.60 ± 0.845 vs. 5.46 ± 0.670 sp/s; one-tailed K-S test, *p* = 0.14). Right, During wall contact at the prefred distance, these same SpVi-TeLC units exhibited significantly elevated firing (20.25 ± 2.263 vs. 16.44 ± 1.640 sp/s; one-tailed K-S test, *p* = 1.64 x 10^-3^). * *p* < 0.0125 -correctd, four comp so s); n.s., not significant.

## STAR* METHODS

### Lead Contact

Further information and requests for resources and reagents should be directed to and will be fulfilled by the Lead Contact, Fan Wang

### Materials Availability

This study did not generate new unique reagents.

## DATA AND SOFTWARE AVAILABILITY

MATLAB scripts and analysis code used in this study will be made available on Github, and processed datasets in NWB (Neurodata Without Borders) format are deposited on DANDI.

## EXPERIMENTAL MODEL DETAILS

Adult C57BL/6J (JAX stock #000664) and vGat-IRES-Cre (JAX stock #016962; Slc32a1^tm2(cre)Lowl/J^) mice were obtained from the Jackson Laboratory. During behavioral and electrophysiological experiments, mice were singly housed under a 12-hour light/dark cycle with ad libitum access to food and water. All experiments were conducted during the light phase. Details regarding the sex and age of all animals are provided in Table S1. All procedures were performed in accordance with protocols approved by the Institutional Animal Care and Use Committees at the Massachusetts Institute of Technology.

## METHOD DETAILS

### Headpost Implantation and Craniotomy

Mice were anesthetized with 3% isoflurane in 0.8 L/min oxygen and positioned in a stereotaxic frame (Model 963, David Kopf Instruments). Anesthesia was maintained at 1.5% isoflurane throughout the surgery. A custom stainless steel headbar (Pololu; 0.060″ ± 0.004″, type 304) and a stainless-steel ground screw were affixed to the skull using C&B Metabond (Parkell, #S380) and reinforced with dental cement (Stoelting). Mice were allowed at least seven days to recover following head-post implantation surgery. On the electrophysiological recording day, a 1 mm × 1 mm craniotomy was created 4.7 mm posterior and 2.0 mm lateral to Bregma using a dental drill (Aseptico). The dura was carefully removed to provide access to the principal trigeminal nucleus (PrV). Craniotomies were acutely sealed with silicone elastomer (Kwik-Cast, WPI).

### Viral Vectors

The AAVs used included AAV-EF1a-DIO-GtACR2-EYFP (Takatoh & Prevosto et al., 2021), AAV-hSyn-FLEX-TeLC-P2A-EYFP-WPRE (Addgene, #135391).

### Viral Injections

For optogenetic silencing experiments, vGat-Cre mice received stereotaxic injections of 300 nl AAV-EF1a-DIO-GtACR2-eYFP into PrV at coordinates 4.7 mm posterior, 2.0 mm lateral to Bregma, and 4.5 mm below the brain surface. For chronic silencing of SpVi neurons projecting to PrV, 300 nl of either AAV-hSyn-FLEX-TeLC-P2A-EYFP-WPRE was stereotaxically injected into caudal SpVi (7.4 mm posterior, 2.0 mm lateral to Bregma, and 5.6 mm below the brain surface).

Injections were performed using beveled glass micropipettes connected to a microsyringe pump (UMP3, WPI) controlled by a Micro4 controller (WPI) at a rate of 30 nl/min. Following injections, craniotomies were sealed with bone wax (Surgical Specialties, #902), and the scalp was closed using nonabsorbable silk sutures (LOOK, #774B).

### Wall-Passing Paradigm

After recovering from headpost implantation, mice were head-fixed using plate clamps (Thorlabs) positioned over a custom linear treadmill (Severson & Xu et al., 2017) to promote running and whisking. Mice were habituated to the treadmill for at least two days, followed by at least three days of habituation on a wall-pass apparatus prior to recording. The passing wall was driven in the mediolateral and anteroposterior dimensions by two linear actuators (Zaber, Models LSM200B-T4 rev 2 and LSM050B-T4 rev 2). During each pass, the wall moved from anterior to posterior across the mouse’s face at a constant velocity of 10 mm/s. In the silicon probe experiments, wall distance from the whisker pad (C2 follicle) was randomly varied across trials between 7 and 23 mm in 2-mm increments (4-mm increments for optogenetic silencing experiments), using a custom MATLAB script. In the tungsten electrode experiments, wall distance was randomly varied between 4 mm and 20 mm in 1 mm increments. All experiments were performed in the dark. After the mediolateral Zaber motor set the wall distance, a randomized delay of 0.1 to 1 s preceded wall movement.

Each recording session in the silicon probe experiments included 15-20 repetitions per wall distance, during which mice freely whisked and interacted with the wall. Tungsten electrode experiments included 10-20 repetitions per distance. Whiskers were illuminated using an infrared LED backlight (Roithner Lasertechnik, #LED660N-66-60) and imaged with a high-speed camera (Basler, Model acA1300-200um) equipped with a telecentric lens (Edmund Optics, #55-349). Prior to each recording, the wall apparatus was aligned to the midline of the mouse’s head, and a calibration image was acquired and converted to physical units (31 pixels/mm). The closest wall distance was defined as the minimum Euclidean distance between the C2 whisker follicle and the wall. Absolute wall distance (in mm) was calculated by adding the calibrated scaling factor to the horizontal position (in 2-mm steps, relative to the closest position) of the stepper motor controlling the wall’s mediolateral position.

### High-Speed Videography

High-speed whisker videos (688 × 688 pixels at 500 Hz) were acquired with a machine-vision camera (Basler acA1300-200um) controlled by custom Python software (CamPy). Illumination was provided by a 940 nm LED (Roithner Lasertechnik) collimated with a condenser lens and diffused to form an even backlight. The whiskers appeared in silhouette against this background and were imaged via a front-surface mirror into a 0.25× telecentric lens (Edmund Optics).

Frame timing was triggered by a Teensy microcontroller, and TTL pulses were recorded alongside electrophysiology to synchronize spikes and video.

### Awake and Anesthetized Wall-Passing Recordings

In a subset of mice (N = 4), high-speed video was continuously acquired during the wall-passing paradigm under both awake and anesthetized conditions. Mice were head-fixed throughout, keeping their position constant relative to the moving wall stimulus. To enable whisker tracking, all whiskers except a single row (C1-C4 and γ or β) were trimmed. While awake, each mouse experienced wall passes with randomized wall distances (7-23 mm in 2-mm increments), repeated for 20 passes per distance. Following the awake session, the same animals were anesthetized via intraperitoneal injection of a fentanyl-based cocktail (0.05 mg/kg fentanyl, 5 mg/kg midazolam, 0.5 mg/kg dexmedetomidine). After a 10-minute induction period, the same wall-passing protocol was repeated under anesthesia. Comparisons between awake and anesthetized conditions were performed within-subject across matched wall distances.

### Electrophysiology Recording: Silicon Probe

Following habituation to the wall-passing paradigm, mice were head-fixed and allowed to run on the treadmill. The craniotomy was exposed and covered with artificial cerebrospinal fluid (aCSF). For wild-type and TeLC experiments, the tip of a 32-channel silicon probe (NeuroNexus, A1×32-Poly2-10mm-50s-177-A32 or A1×32-Poly3-10mm-50s-177) was coated with the fluorescent dye Dil (Invitrogen™, #V22885). For optogenetic silencing experiments, a 473 nm laser (Cobolt, 06-MLD 473) was connected to a ferrule attached to a 32-channel silicon optrode (NeuroNexus, A1×32-Poly2-10mm-50s-413, 105 µm/125 µm, 0.22 NA). The probe or optrode was advanced approximately 5 mm into the brain using a micromanipulator (Sutter Instruments, Model MPC-325) until it reached ventral PrV and well-isolated units responsive to manual whisker stimulation were encountered. The brain was allowed to relax for at least 40 minutes to ensure stable recordings. Neural signals were hardware-filtered on the headstage (Blackrock Microsystems, Cereplex M, #10532) between 0.3 Hz and 7.5 kHz for spikes and digitized at 30 kHz using Cereplex Direct software (Blackrock Microsystems, Version 7.0.5). Analog voltage signals from the anterior-posterior stepper motor were simultaneously acquired at 30 kHz. Wall pass start and end times were determined from the rising and falling edges of these voltage signals to synchronize electrophysiological recordings with wall movement. Recording sites were subsequently confirmed by post hoc histological analysis.

### Electrophysiology Recording: Single Tungsten Electrode

One day prior to recording, whiskers and facial fur on the left side of the face were trimmed using fine forceps and microdissection scissors (FST, #91500-09) under isoflurane anesthesia (1.5%), sparing the C-row whiskers (γ or β, C1–C5). On the recording day, mice were head-fixed and allowed to run on the treadmill. The craniotomy was exposed and kept covered with aCSF. The tip of a tungsten microelectrode (0.005″ diameter, 5 MΩ, 8° tip angle; A-M Systems, #575300) was coated with the fluorescent dye Dil and lowered approximately 5 mm into the brain using a micromanipulator until it reached ventral PrV. Electrical activity was monitored via audio output from the amplifier (WPI, DAM80). Manual whisker stimulation was performed to identify a well-isolated, whisker-responsive unit. The brain was allowed to relax for at least 20 minutes to achieve stable recordings. Differential signals between the recording and reference electrodes were amplified 10,000× and bandpass filtered between 300 and 3,000 Hz (WPI, DAM80). Following recording, receptive fields and response types (rapidly adapting [RA] or slowly adapting [SA]) were manually classified. Analog voltage signals from the amplifier and the anterior-posterior stepper motor were acquired using Cereplex Direct at a sampling rate of 30 kHz. Wall pass start and end times were extracted from the rising and falling edges of the stepper motor TTL signals to synchronize electrophysiological recordings with wall movement. All recording sites were confirmed by post hoc histological analysis.

### Isolation of Single Unit Activity

For silicon probe data, initial single-unit isolation (spike sorting) was performed using Kilosort2.5 (Pachitariu et al., 2016), followed by manual curation in Phy (github.com/cortex-lab/phy; Rossant, 2025). Spike times and waveforms for isolated clusters were exported from Phy. Most units exhibited narrow spike widths, defined by a trough-to-peak latency of less than 0.4 ms. Duplicate units, identified as those with high Pearson’s correlation coefficients (> 0.1) in spike counts binned at 1 ms resolution, were excluded from further analysis. For tungsten electrode data, spike waveforms were extracted by thresholding high-pass filtered traces (500 Hz) and clustered using MClust-4.4 (A.D. Redish, http://redishlab.neuroscience.umn.edu/MClust/MClust.html).

Isolation quality was assessed using the following metrics: signal-to-noise ratio (SNR), defined as the mean waveform amplitude divided by the standard deviation (SD) of waveform noise; and false alarm (FA) rate, defined as the percentage of spikes with inter-spike intervals (ISI) shorter than 0.8 ms. Units with an SNR less than 5 SDs or an FA rate greater than 2% did not meet single-unit isolation quality criteria and were excluded from subsequent analyses. For silicon probe data, in wild-type mice, 1141 of 1358 units (36 mice), in TeLC experiments, 587 of 665 units (11 mice), and for Tungsten recordings (46 of 46 units (14 mice), passed isolation quality and minimum firing rate (>1 Hz) criteria and were subsequently entered into population analyses. For acute whisker cutting experiments, only units demonstrating stable waveform properties both before and after cutting were included in analyses.

### Acute Whisker Cutting Manipulation

During recording sessions involving acute whisker cutting, neural responses were first recorded while the full whisker array was intact. Each session included 15 to 20 repetitions of wall passes over approximately one hour. To best guess the primary whisker to spare during acute trimming, the receptive fields of PrV neurons were mapped by manually stimulating individual whiskers with a cotton-tipped applicator while monitoring spike responses. All whiskers except the designated spared whisker were then cut down to the base of the shaft using microdissection scissors (FST, Cat# 91500-09). After trimming, the spared whisker and surrounding whisker pad were brushed with a cotton-tipped applicator to stimulate all touch-sensitive recorded units. The wall-passing procedure was then repeated for an additional 15 to 20 trials. After completing the post-cutting wall passes, the whisker and pad were brushed once more to confirm the continued presence of responsive units. Electrophysiological signals were continuously recorded throughout the pre-cut, cutting, brushing, and post-cut stimulation periods.

### Optogenetic Silencing of PrV Local Interneurons

Light pulses were generated using a pulse generator (Doric, #OTPG-4), with laser timing synchronized to wall movement via Doric software. Continuous 473 nm light stimulation (2 mW) was delivered by a laser (Cobolt, #06-MLD 473) throughout each wall pass, with a 5-second inter-trial interval. Each recording session consisted of 30 to 40 repetitions at each wall distance, ranging from 7 to 23 mm in 4 mm increments, conducted over approximately two hours with randomized “Laser Off” and “Laser On” trials.

### Histology

Following recordings, mice were deeply anesthetized and transcardially perfused with phosphate-buffered saline (PBS), followed by 4% paraformaldehyde in PBS. Brains were post-fixed overnight and then cryoprotected in 30% sucrose in PBS at 4°C. Tissue was embedded in OCT compound (Sakura Finetek) and coronally sectioned at 100 μm using a cryostat (Leica Biosystems). For wild-type, optogenetic inhibition, and tungsten recording experiments, sections containing PrV were stained with Neurotrace Blue (1:500; Thermo Fisher Scientific, #N21479) in 0.3% Triton X-100/PBS. For TeLC experiments, sections containing PrV and SpVi underwent immunostaining with goat anti-GFP (1:500; Rockland, #600-101-215) in blocking buffer, followed by either Alexa Fluor 488 (1:500; Jackson ImmunoResearch, #705-545-147) or Alexa Fluor 647 (1:500; Jackson ImmunoResearch, #705-606-147) donkey anti-goat secondary antibodies and Neurotrace Blue (1:500; Thermo Fisher Scientific, #N21479) in blocking buffer. The sections were then briefly rinsed with 0.3% Triton X-100/PBS and mounted on slides with a custom-made mounting medium. All histological images were acquired with either a Keyence microscope (BZ-X800) or a Zeiss LSM700 confocal microscope.

## QUANTIFICATION AND STATISTICAL ANALYSIS

All analyses were performed in MATLAB (MathWorks, versions 2021b and 2023b). Data are reported as mean ± standard errors of the mean (SEM) unless otherwise noted. Measures of central tendency, variability, test statistics, p-values, degree of freedom, and figure panel assignments for all analyses are detailed in Table S6. We analyzed single-unit electrophysiological recordings from n = 1,187 units in PrV across 63 mice, as well as high-speed video recordings from four sessions in four mice. The video data comprised 15,679,458 frames, totaling 522.6 minutes. Sample sizes were not predetermined.

### Video Analysis

The backbone of each whisker was tracked at subpixel-resolution using the Janelia Whisker Tracker (Clack et al., 2012), yielding a set of “trajectories” (tracked objects in image X-Y coordinates) for each frame. All subsequent processing of whisker angle (*θ*) was conducted in MATLAB according to published methods (Pammer & O’Connor et al., 2013; Severson & Xu et al., 2017), with several modifications described below in “Video analysis” subsections.

### Video Analysis: Identifying Tracked Whiskers

To identify the same whiskers across frames (linking), whiskers were ordered by follicle x-position. Trajectories with follicle x- or y-position outside the region of interest (bounding box), with arc length shorter than 150 pixels (4.65 mm), or whisker “score” (Janelia whisker tracker measurements) less than 600 were ignored. Post-hoc median filtering (3-frame window) was applied to smooth over single-frame linking errors.

### Video Analysis: Face Masking

Computing time series of whisker angle relies on being able to reliably track the base of the whisker near the follicle location across all video frames. As previously described (Pammer & O’Connor et al., 2013; Severson & Xu et al., 2017), we used a dynamic “mask” to truncate the tracked whisker trajectories near the face to remove “noise” due to occlusion by facial fur. The follicle location was estimated by extrapolating past the intersection of the whisker and the mask using the angle at the whisker base (Pammer & O’Connor et al., 2013). We fit a separate mask for each frame, obtained using a custom algorithm that fitted a smoothing spline to the contour of the face. We blurred the image using a Gaussian filter, binarized the image, and detected the contour by taking a spatial derivative along the y-dimension. The pixel locations of the facial contour were thus detected as the intersection between the white background and black shadow cast by the face. A smoothing spline was then fitted to the facial contour positions and shifted upward by ∼10 pixels to yield a 20-point facemask that intersected with the whisker trajectories. Frames in which a tracked whisker did not intersect the facemask were considered missing data.

### Data Analysis: Wall-Induced Whisker Retraction, Awake vs Anesthetized

For each of the tracked whiskers (C1, C2, and Greek), whisker angle (*θ*) was measured relative to the mediolateral axis of the face. The midpoint of the whisker angle trace (*θ_mid_*) was defined as the slowly varying centerline of *θ*. To obtain *θ_mid_*, *θ* was first smoothed using a Savitsky-Golay filter (21 frame, third order), followed by interpolation of the peaks and valleys to obtain the upper and lower envelope of the *θ* trace. *θ_mid_* was then calculated as the mean of this envelope. For each wall pass, *θ_mid_* was computed during both the baseline period (−4.5 to 0 s, with t=0 marking wall onset) and the wall-passing period (3 to 11 s). Baseline-subtracted *θ_mid_* (Δ*θ_mid_*) was then calculated for each pass. For each mouse (N = 4 mice), Δ*θ_mid_* was averaged across wall distances and repetitions to obtain a single value per whisker and condition (awake or anesthetized). Data were aggregated across mice into a summary table (mouse × whisker × condition), and paired comparisons were made between awake and anesthetized conditions for each whisker.

To compare the effect of wall-induced whisker retraction of awake versus anesthetized conditions, we performed two-tailed paired t-tests across mice (within-subjects) for each whisker. The null hypothesis was that mean Δ*θ_mid_* was equal in awake and anesthetized conditions. Effect size was computed as the mean difference between conditions (awake minus anesthetized), and Bonferroni correction was applied to control for multiple comparisons across whiskers (*α* =0.0167).

### Data Analysis: Spike Density Function Calculation

Spike density functions (SDFs) were computed by aligning spike times to the onset of wall-pass trials, with time zero defined as the moment when the wall began moving toward the mouse. For all trial conditions, two epochs were defined consistently: a baseline period from −4.5 to 0 s, and a wall-pass period from 3 to 11 s. Spike rasters were generated from the aligned spike times using 1 ms bins. SDF curves were then produced by convolving these spike rasters with a Gaussian filter (5 ms SD, full kernel). SEMs were calculated as the SD of the SDF across trials divided by the square root of the number of trials in each condition. The resulting SDFs and SEMs were averaged across repetitions of the same wall distance and stored for each cell.

### Data Analysis: Wall Distance Tuning Calculation

Wall distance tuning curves were generated by uniformly binning distance measurements into nine 2-mm-wide bins with centers at 7, 9, 11, 13, 15, 17, 19, 21, and 23 mm. Distances falling outside the bin edges were excluded. Peri-event time histograms (PETHs) were computed by binning instantaneous firing rates in 200-ms intervals, aligned to the onset of anterior-posterior wall movement as defined by the stepper motor TTL pulse (t=0). Each wall pass trial lasted approximately 14 seconds. Average firing rates during the wall pass period (3–11 s) were computed from the PETH and compared to baseline firing rates calculated from −4.5 to 0 s. For each distance, mean firing rates were averaged across repeated passes to construct the tuning curve. Standard errors were calculated by dividing the SD across repetitions by the square root of the number of repetitions for each distance. CIs were estimated using bootstrap resampling (10,000 iterations) and defined as the 95th percentile range. Z-scored tuning curves were computed by subtracting the mean from firing rate across all baseline periods in the recording session and dividing by the corresponding SD. Uniform binning enabled concatenation of data across sessions for population-level analyses, including decoding analyses described later. Sessions with fewer than nine presented distances were excluded from population analyses. For population heatmaps, each neuron’s tuning curve was normalized to its own minimum and maximum firing rates, and the color scale represents the resulting normalized mean spike rate.

### Data Analyses: Characterizing Single Unit Responses

To assess responsiveness to wall stimulation, one-tailed Wilcoxon signed-rank tests (*p* < 0.05) were performed on each of 1,187 units. A unit was classified as “wall-activated” if its mean firing rate during the wall pass period (3-11 s) at any distance was significantly higher than its baseline firing rate (−4.5-0 s). Conversely, a unit was classified as “wall-suppressed” if its mean firing rate during the wall pass period at any distance was significantly lower than baseline. Units exhibiting significant increases at some distances and decreases at others were classified as both “wall-activated” and “wall-suppressed.”

Wall distance tuning was assessed using two-tailed Wilcoxon signed-rank tests across all pairs of distances. A unit was classified as “distance-tuned” if the mean firing rate at one or more distances significantly differed from that at other distances (*p* < 0.05, corrected for multiple comparisons using the Benjamini-Hochberg false discovery rate method).

For comparisons of baseline firing rate distributions between distance-tuned and untuned units, the mean and 95% confidence intervals (CI) were calculated using bootstrap resampling (10,000 iterations). The CI was determined by calculating half the range between the upper and lower bounds of the bootstrap interval. Results were reported as mean ± CI. Statistical differences between distributions were assessed using two-sample Kolmogorov-Smirnov (K-S) tests (‘kstest2’ function in MATLAB), performed with one- or two-tailed hypotheses as dictated by the experimental design (see Table S6). Significance was determined at *p* < 0.05, with Bonferroni correction for multiple comparisons. The same procedures were applied for comparing average firing rates during wall passes and baseline periods between distance-tuned and untuned populations.

To further characterize tuning profile within “wall-activated” and “distance-tuned” units, we performed one-tailed Wilcoxon signed-rank tests comparing mean firing rates at each non-closest distance (≥ 9 mm) to the closest distance (7 mm). Units with significantly higher firing rates at any non-closest distance were classified as “map units,” while those exhibiting maximal firing at the closest distance (7 mm) were classified as “proximity units.” Units that met criteria for “wall-activated” and “distance-tuned” but did not fulfill the criteria for either map or proximity were labeled “ambiguous” and excluded from decoding and population cross-validation analyses.

For analyses of acute whisker cutting experiments, mean firing rates during baseline and wall-pass periods were compared between full whisker array (control) and single-whisker (cut) conditions. Pairwise comparisons of firing rates were conducted using two-tailed Wilcoxon signed-rank tests. To assess individual neuron responses to whisker cutting, the proportions of units exhibiting significant increases, decreases, or no changes in firing rates were computed. Significance thresholds were corrected for multiple comparisons using Bonferroni adjustment. To evaluate time-dependent plasticity, firing rates for each unit were partitioned into three temporal segments: the first half of wall-pass trials (full whisker array), the second half (full whisker array), and the post-manipulation period. For each unit, mean firing rates were computed separately for these periods. Adaptation was assessed by comparing firing rates between the first and second halves using two-tailed Wilcoxon signed-rank tests. A non-significant result (*p* > 0.05) indicated stable firing prior to whisker cutting. Firing rates from the second half of the full whisker array period were then compared to post-cutting periods using two-sided Wilcoxon signed-rank tests, where significant differences indicated effects attributable to whisker cutting.

To compare the proportions of map, proximity, ambiguous, suppressed, or untuned cells between SpVi-TeLC mice and wild-type mice, a one-tailed Z-test for proportions was performed. The null hypothesis (*H*₀) stated that the proportion of each cell type in the SpVi-TeLC population (*p₂*) was less than or equal to that in the wild-type population (*p₁*), while the alternative hypothesis (*Hₐ*) posited that *p₂* > *p₁*. Proportions (*p₁* & *p₂*) were calculated as the number of cells of a given type (*x₁* & *x₂*) divided by the total number of recorded units (*n₁* and *n₂*) in each group. A pooled proportion (*p_pooled_*) was used to estimate the standard error (SE) of the difference between groups:

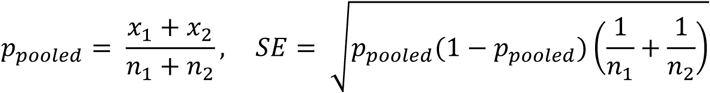

The Z-statistic was computed as:

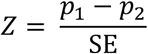

P-values were calculated using MATLAB’s cumulative normal distribution function (‘normcdf’ in MATLAB) and evaluated at *α* = 0.05. A p-value below this threshold indicated a significantly higher proportion of the specified cell type in SpVi-TeLC mice compared to wild-type mice.

To compare the firing rate distributions between SpVi-TeLC and wild-type populations (Group 1 and Group 2), we computed the mean and 95% CI for each group using bootstrap resampling (10,000 iterations). For each group, the results were reported as mean ± CI. To assess the statistical significance of differences between the distributions of the two groups, we performed a two-sample KS test. The test was conducted with a specified directional hypothesis (one- or two-tailed, depending on the experimental design), and the resulting p-value was reported to determine the significance (*p* < 0.05, with Bonferroni correction for multiple comparisons) of the observed differences.

### Data Analysis: Decoding of Wall Distance from Population or Single Neurons Activity

For population decoding of wall distance, we applied a population decoder (Neural Decoding Toolbox, www.readout.info; Meyers, 2013; Vincis & Chen et al., 2020). Units from sessions with all nine distances presented that passed spike sorting quality metrics (1,141 total units) were concatenated into a pseudo-population. Populations were subsampled into sub-populations of equal size (118 units), containing units designated as either “map”, “proximity”, or “untuned” cells. Units in the “shuffled” subpopulation were randomly sampled from map and proximity subpopulations and had their trial distance labels shuffled with replacement. Spike timestamps for each unit were aligned to the onset of anterior-posterior wall movement (time 0) and were binned using a 50 ms bin size with a 150 ms sliding window to create a firing rate matrix, where each row corresponds to a trial and each column corresponds to a time bin. The firing rate matrix spans from time 0 to 14 seconds. To ensure equal trial numbers across conditions (i.e., wall distances), trials were subsampled five times to match 15 trials per condition. The binned data and their corresponding condition labels were randomly split into 10 subsets—9 used for training the classifier and 1 reserved for validation. The process was repeated 10 times for each unique partition for 10-fold cross-validation to assess decoding accuracy. The classifier operates by learning a mean population vector (template) for each class from the training set. During testing, it calculates the Pearson correlation coefficient between a test point and the learned templates. During validation and testing, the class with the highest correlation value is assigned as the predicted label. Decoding accuracy (true positive rate) was averaged over the 3-11 second wall-pass period to obtain the overall decoding accuracy for the sampling epoch. In addition to the decoding accuracy, confusion matrices were computed at each time step. To assess variability, the entire subsampling procedure was repeated 10 times, and standard error was calculated by computing the SD of decoding accuracy across runs and dividing by the square root of 10.

For single-neuron decoders, we quantified how well individual neurons discriminated wall distance. For each unit, spike trains were grouped according to nine discrete wall distances (7, 9, 11, 13, 15, 17, 19, 21, and 23 mm). For each distance, spike times were aligned to trial onset and converted into spike count vectors using 200 ms non-overlapping bins spanning the 3–11 s wall contact period. These binned firing rate profiles were used to construct condition-specific templates for distance classification. Decoding was performed using a Euclidean distance classifier, which assigned each test trial to the distance condition whose mean firing-rate vector (template) was closest in Euclidean space. Cross-validation was implemented as a leave-two-out procedure (xval = 2), in which two trials per condition were held out as test data, and the remaining trials were used for training. This was repeated exhaustively across all combinations of held-out pairs, such that every trial pair served as a test set at least once. Classification accuracy was computed as the fraction of correctly identified test trials, averaged across all cross-validation iterations. For shuffled controls, we performed a trial-shuffled permutation in which class labels were randomly reassigned across conditions (n = 1000 iterations).

For both population and single-neuron decoders, differences in mean decoding accuracy and error across wall distances were assessed using Wilcoxon rank-sum tests. Separate tests were performed for each of the nine wall distances, and p-values were corrected for multiple comparisons using BH-FDR to compare proximity and map populations.

### Data Analysis: Acute Whisker Cutting Manipulation

To test for the effect for acute whisker cutting on firing rates, we computed each unit’s mean firing rate during baseline and wall-passing periods on each repetition for the full whisker array (control) or single-whisker condition (manipulation). Pairwise comparison of mean firing rate for neural population and individual units between conditions was performed via two-sided Wilcoxon signed-rank test. To evaluate how individual neurons responded to whisker cutting, we computed the proportion of units that exhibited significant increases, decreases, or no change in firing rate between conditions. P-values were corrected for multiple comparisons across units using the BH-FDR procedure.

### Data Analysis: Modulation Index in Acute Optogenetic Experiments

We quantified the acute optogenetic effects on firing rates using a modulation index (MI) computed for each unit as:

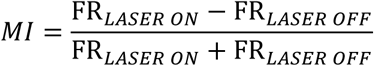

where FR_LASER ON_ and FR_LASER OFF_ denote the mean firing rates during matched laser-on and laser-off wall-passing periods, respectively. For each unit, we obtained a bootstrap CI on the MI and classified the unit as significantly modulated if the CI did not include zero. Significantly modulated units were further assigned to one of three categories based on their MI: strongly inhibited (MI ≤ −0.5), inhibited (−0.5 < MI < 0), or disinhibited (MI > 0). Units whose MI confidence intervals overlapped zero were classified as no change. All analyses were restricted to units that passed our unit inclusion criteria (see Isolation of Single Unit Activity).

### Computational Simulation of Brainstem Circuitry

To model the tuning properties of PrV neurons, we simulated a pool of TG afferents whose activities decreased monotonically with wall distance, consistent with their mechanical sensitivity. These proximity-based TG inputs projected to PrV neurons via a direct excitatory (lemniscal) pathway and an indirect inhibitory pathway (SpVi inhibitory projections to PrV). TG tuning curves were modeled as sigmoidal functions with thresholds spanning the distance range. The activity of each PrV neuron 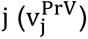 in response to wall distance x ∈ [0,1] was defined as:

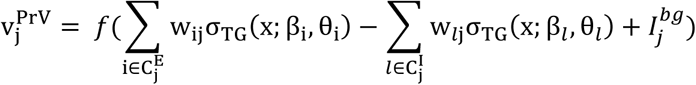

where

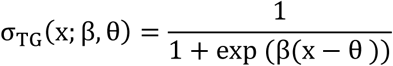

is the TG neuron’s sigmoidal tuning function. Here, β and θ control sensitivity and selectivity to wall distance, respectively. *f*(*x*) = max {*x*, 0} is the non-linear activation function for PrV neurons and 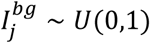 is the background excitatory input to PrV neuron *j*. 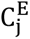denotes the set of TG neurons providing direct excitatory input to PrV neuron j, with synaptic weight 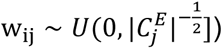; 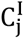 denotes TG neurons projecting indirectly to PrV neuron *j* via inhibitory neurons in SpVi, with weight 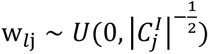. Subscript *i* and *l* denote each TG neuron in set 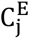 and set 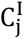. For simplicity, inhibitory neurons were modeled as providing a sign-inverted copy of TG activity, such that 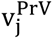 reflect a linear combination of excitatory and inhibitory streams.

Intuitively, the subtraction of two sigmoidal functions can yield a non-monotonic, Gaussian-like function. Here we generalize this by considering a pool of sigmoidal functions (TG neurons). Specifically, we implemented several assumptions:

(1) Sparse connectivity: Both 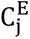 and 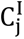 represented small subsets of TG neurons. Parameter β_i_, θ_i_, β_*l*_, θ_*l*_, w_ij_, w_*l*j_ were independently sampled for each PrV neuron. The number of inhibitory inputs 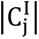 per PrV neuron *j* was treated as a random variable sampling from an exponential distribution parametrized by λ_*c*_ (truncated to the interval [1,30] to preserve E/I balance):

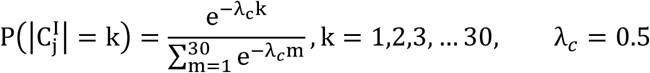 The number of excitatory inputs was fixed at 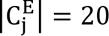.
(2) Skewed distribution of TG tuning. The TG threshold parameter θ was drawn from a truncated exponential distribution within [0,1]:

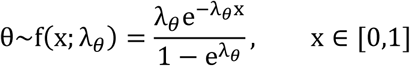 where λ_*θ*_ describes the skewness of the distribution. This skewed distribution increases the density of high-threshold TG neurons that only fire at proximal distances, thereby enhancing encoding resolution and reducing overall metabolic cost (efficient coding). We used λ_*θ*_ = 2 for the excitatory inputs and λ_*θ*_ = 4 for the more skewed inhibitory pathway. Sensitivity parameters were sampled as β_i_, β_*l*_∼N(20, 5). Untuned neurons were modeled by explicitly weakening all weights for a fraction of neurons to 1% (i.e., p(untuned) = 0.4).

The input (distance) range x was normalized to [0,1] for simplicity. We sampled 99 uniformly spaced points corresponding to wall-distance presentations. To evaluate tuning curves, we averaged simulated activity into nine discrete distance bins [0,…, 8]. All neurons with flat tuning curves (max – min < 0.1) were considered untuned. Neurons with max activity below zero were considered suppressed. Among the remainder, proximity cells were defined as neurons with maximal responses at x = 0, while map cells were defined as neurons with peak responses at intermediate or far distances (x > 0). In total, we modeled 1,225 PrV neurons.

To simulate partial silencing of inhibitory projections, the distribution of inhibitory inputs was truncated to approximately 50% of its original range [1,15], such that 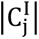 for each PrV neuron is decreased by:

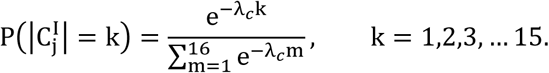

---

**Table S1.** Mouse Metadata and Figure Appearances.

| Mouse ID | Sex | Mouse Line | Birth Date | Recording Dates | Experiments | Appearances - Main Figures |  |  |  |  |  |
| --- | --- | --- | --- | --- | --- | --- | --- | --- | --- | --- | --- |
|  |  |  |  |  |  | 1 | 2 | 3 | 4 | 5 | 6 |
| Ephys1 | M | C57BL/6J | 11/08/22 | 02/15/23-02/17/23 | PrV Si probe, WT |  | D-F | B-D,F |  | C-E |  |
| Ephys2 | M | C57BL/6J | 11/08/22 | 02/08/23-02/10/23 | PrV Si probe, WT | E | A-F | B-D,F |  | C-E |  |
| Ephys3 | M | C57BL/6J | 09/20/22 | 02/01/23-02/03/23 | PrV Si probe, WT |  | D-F | B-D,F |  | C-E |  |
| Ephys5 | M | C57BL/6J | 06/22/21 | 08/19/21-08/24/21 | PrV Tungsten |  |  |  |  |  |  |
| Ephys6 | M | C57BL/6J | 06/22/21 | 09/17/21-09/20/21 | PrV Tungsten |  |  |  |  |  |  |
| Ephys7 | M | C57BL/6J | 06/22/21 | 09/13/21-09/14/21 | PrV Tungsten |  |  |  |  |  |  |
| Ephys8 | F | C57BL/6J | 01/25/22 | 06/15/22 | PrV Tungsten |  |  |  |  |  |  |
| Ephys9 | M | C57BL/6J | 08/24/21 | 10/15/21-10/19/21 | PrV Tungsten |  |  |  |  |  |  |
| Ephys12 | M | C57BL/6J | 05/14/22 | 08/08/22 | PrV Tungsten |  |  |  |  |  |  |
| Ephys13 | F | C57BL/6J | 04/12/22 | 06/29/22-07/06/22 | PrV Tungsten |  |  |  |  |  |  |
| Ephys14 | F | C57BL/6J | 04/12/22 | 07/27/22-08/04/22 | PrV Tungsten |  |  |  |  |  |  |
| Ephys15 | M | C57BL/6J | 05/14/22 | 08/11/22-08/25/22 | PrV Tungsten |  |  |  |  |  |  |
| Ephys16 | M | C57BL/6J | 05/14/22 | 09/01/22-09/06/22 | PrV Tungsten |  |  |  |  |  |  |
| Ephys17 | M | C57BL/6J | 05/14/22 | 09/09/22 | PrV Tungsten |  |  |  |  |  |  |
| Ephys18 | M | C57BL/6J | 07/12/22 | 09/29/22-10/04/22 | PrV Tungsten |  |  |  |  |  |  |
| Ephys19 | M | C57BL/6J | 07/12/22 | 10/17/22 | PrV Tungsten |  |  |  |  |  |  |
| Ephys20 | M | C57BL/6J | 07/12/22 | 10/27/22-11/01/22 | PrV Tungsten |  |  |  |  |  |  |
| Ephys24 | F | C57BL/6J | 10/25/22 | 01/31/23 | PrV Si probe, WT |  | D-F | B-D,F |  | C-E |  |
| Ephys25 | M | C57BL/6J | 11/08/22 | 02/22/23-02/25/23 | PrV Si probe, WT |  | D-F | B-D,F |  | C-E |  |
| Ephys26 | M | C57BL/6J | 11/08/22 | 02/26/23-02/28/23 | PrV Si probe, WT |  | D-F | B-D,F |  | C-E |  |
| Ephys27 | F | C57BL/6J | 11/15/22 | 03/21/23-03/23/23 | PrV Si probe, WT |  | D-F | B-D,F |  | C-E |  |
| Ephys28 | F | C57BL/6J | 11/15/22 | 03/24/23-03/27/23 | PrV Si probe, WT |  | D-F | B-D,F |  | C-E |  |
| Ephys29 | F | C57BL/6J | 11/29/22 | 03/28/23-03/30/23 | PrV Si probe, WT |  | D-F | B-D,F |  | C-E |  |
| Ephys30 | M | C57BL/6J | 02/27/23 | 05/24/23 | PrV Si probe, WT |  | D-F | B-D,F |  | C-E |  |
| Ephys32 | F | C57BL/6J | 03/14/23 | 06/15/23 | PrV Si probe, WT |  | D-F | B-D,F |  | C-E |  |
| Ephys33 | F | C57BL/6J | 03/14/23 | 06/02/23 | PrV Si probe, WT |  | D-F | B-D,F |  | C-E |  |
| Ephys34 | M | C57BL/6J | 02/27/23 | 07/11/23 | PrV Si probe, WT |  | D-F | B-D,F |  | C-E |  |
| Ephys36 | F | C57BL/6J | 03/14/23 | 06/08/23 | PrV Si probe, WT | D | D-F | B-D,F |  | C-E |  |
| Ephys40 | M | C57BL/6J | 03/28/23 | 06/26/23 | PrV Si probe, WT |  | A-F | B-D,F |  | C-E |  |
| Ephys41 | M | C57BL/6J | 03/28/23 | 06/27/23 | PrV Si probe, WT |  | D-F | B-D,F |  | C-E |  |
| Ephys42 | F | C57BL/6J | 05/23/23 | 07/25/23 | PrV Si probe, WT |  | D-F | B-D,F |  | C-E |  |
| beh104 | M | C57BL/6J | 06/06/23 | 12/04/23 | whisker behavior |  |  |  |  |  |  |
| beh101 | F | C57BL/6J | 09/26/23 | 12/12/23 | whisker behavior |  |  |  |  |  |  |
| beh102 | M | C57BL/6J | 09/26/23 | 12/13/23 | whisker behavior |  |  |  |  |  |  |
| beh103 | F | C57BL/6J | 08/15/23 | 12/12/23 | whisker behavior |  |  |  |  |  |  |
| Ephys72 | M | vGAT-ires-Cre | 11/27/23 | 03/07/23-03/08/23 | PrV optrode local GtACR |  |  |  |  |  |  |
| Ephys73 | F | vGAT-ires-Cre | 11/27/23 | 03/11/23-03/12/23 | PrV optrode local GtACR |  |  |  |  |  |  |
| Ephys75 | M | C57BL/6J | 04/02/24 | 05/22/24 | PrV Si probe, WT + 1WC | C | D-F | B-D,F | B,K | C-E |  |
| Ephys76 | F | C57BL/6J | 04/02/24 | 06/26/24 | PrV Si probe, WT + 1WC |  | D-F | B-D,F | B-F,K | C-E |  |
| Ephys77 | F | C57BL/6J | 04/22/24 | 08/14/24 | PrV Si probe, WT + 1WC |  | D-F | B-D,F | B | C-E |  |
| Ephys78 | F | C57BL/6J | 04/02/24 | 07/02/24 | PrV Si probe, WT + 1WC |  | D-F | B-D,F | B | C-E |  |
| Ephys79 | M | C57BL/6J | 04/02/24 | 06/24/24 | PrV Si probe, WT + 1WC |  | D-F | B-D,F | A-D,K | C-E |  |
| Ephys81 | F | C57BL/6J | 04/02/24 | 07/03/24 | PrV Si probe, WT + 1WC |  | A-F | B-D,F | B,G,H,K | C-E |  |
| Ephys83 | M | vGAT-ires-Cre | 03/09/24 | 07/09/24 | PrV Si probe, TeLC in SpVi |  |  |  |  | C-E |  |
| Ephys85 | M | vGAT-ires-Cre | 03/09/24 | 07/11/24 | PrV Si probe, TeLC in SpVi |  |  |  |  | C-E |  |
| Ephys87 | F | C57BL/6J | 03/12/24 | 07/24/24 | PrV Si probe, WT + 1WC |  | D-F | B-D,F | B | C-E |  |
| Ephys88 | M | C57BL/6J | 05/14/24 | 09/05/24 | PrV Si probe, WT + 1WC |  | A-F | B-D,F | B,I,J,K | C-E |  |
| Ephys89 | F | C57BL/6J | 04/22/24 | 07/26/24 | PrV Si probe, WT + 1WC |  | D-F | B-D,F | B,K | C-E |  |
| Ephys92 | F | C57BL/6J | 04/22/24 | 08/15/24 | PrV Si probe, WT + 1WC |  | D-F | B-D,F | B,K | C-E |  |
| Ephys94 | M | vGAT-ires-Cre | 05/22/24 | 08/17/24 | PrV Si probe, TeLC in SpVi |  |  |  |  | C-E |  |
| Ephys96 | F | vGAT-ires-Cre | 05/18/24 | 08/20/24 | PrV Si probe, TeLC in SpVi |  |  |  |  | C-E |  |
| Ephys97 | F | vGAT-ires-Cre | 03/01/24 | 11/14/24 | PrV Si probe, TeLC in SpVi |  |  |  |  | C-E |  |
| Ephys100 | M | vGAT-ires-Cre | 06/20/24 | 09/08/24 | PrV Si probe, TeLC in SpVi |  |  |  |  | C-E |  |
| Ephys101 | M | C57BL/6J | 05/14/24 | 09/09/24 | PrV Si probe, WT + 1WC |  | D-F | B-D,F | B,K | C-E |  |
| Ephys102 | F | vGAT-ires-Cre | 06/20/24 | 09/12/24 | PrV Si probe, TeLC in SpVi |  |  |  |  | C-E |  |
| **Ephys105** | M | *C57BL/6J* | 07/02/24 | 09/23/24 | PrV Si probe, WT + 1WC |  | D-F | B-D,F | B | C-E |  |
| **Ephys106** | M | *C57BL/6J* | 07/02/24 | 09/24/24 | PrV Si probe, WT + 1WC |  | A-F | B-D,F | B,I,J,K | C-E |  |
| **Ephys107** | M | *C57BL/6J* | 07/02/24 | 09/25/24 | PrV Si probe, WT + 1WC |  | D-F | B-D,F | B,K | C-E |  |
| **Ephys108** | M | *C57BL/6J* | 07/02/24 | 09/26/24 | PrV Si probe, WT + 1WC |  | D-F | B-D,F | B,E,F,K | C-E |  |
| **Ephys114** | F | *vGAT-ires-Cre* | 05/22/24 | 11/18/24 | PrV Si probe, TeLC in SpVi |  |  |  |  | C-E |  |
| **Ephys119** | M | *vGAT-ires-Cre* | 08/20/24 | 12/08/24-12/09/24 | PrV Si probe, TeLC in SpVi |  |  |  |  | B-E |  |
| **Ephys121** | F | *vGAT-ires-Cre* | 08/20/24 | 12/14/24-12/15/24 | PrV Si probe, TeLC in SpVi |  |  |  |  | C-E |  |
| **Ephys122** | M | *vGAT-ires-Cre* | 09/09/24 | 12/16/24-12/17/24 | PrV Si probe, TeLC in SpVi |  |  |  |  | C-E |  |
| **Ephys126** | M | *C57BL/6J* | 12/03/24 | 03/20/25-03/21/25 | PrV Si probe, WT + 1WC |  | D-F | B-D,F | B | C-E |  |
| **Ephys127** | F | *C57BL/6J* | 11/03/24 | 03/24/25-03/25/25 | PrV Si probe, WT + 1WC |  | D-F | B-D,F | B,K | C-E |  |
| **Ephys128** | F | *C57BL/6J* | 12/31/24 | 03/26/25-03/27/25 | PrV Si probe, WT + 1WC |  | D-F | B-D,F | B,K | C-E |  |

|  | Appearances - Supplemental Figures |  |  |  |  |  |  |
| --- | --- | --- | --- | --- | --- | --- | --- |
| Mouse ID | S1 | S2 | S3 | S4 | S5 | S6 | Other Notes |
| Ephys1 |  |  | B-E | A-D | B,C | H,I |  |
| Ephys2 |  |  | A-E | A-E | A-C | H,I | Unit 5 in 20230208_s1 is 1st example in Fig. 2A-C, S3A, S4E, S5A |
| Ephys3 |  |  | B-E | A-D | B,C | H,I |  |
| Ephys5 |  | F | J-M |  |  |  |  |
| Ephys6 |  | F | J-M |  |  |  |  |
| Ephys7 |  | F | J-M |  |  |  |  |
| Ephys8 |  | B,F | J-M |  |  |  |  |
| Ephys9 |  | C-F | J-M |  |  |  |  |
| Ephys12 |  | F | J-M |  |  |  |  |
| Ephys13 |  | D-F | J-M |  |  |  |  |
| Ephys14 |  | F | J-M |  |  |  |  |
| Ephys15 |  | F | J-M |  |  |  |  |
| Ephys16 |  | F | J-M |  |  |  |  |
| Ephys17 |  | F | J-M |  |  |  |  |
| Ephys18 |  | F | J-M |  |  |  |  |
| Ephys19 |  | F | J-M |  |  |  |  |
| Ephys20 |  | F | J-M |  |  |  |  |
| Ephys24 |  |  | B-E | A,C | B,C | H,I |  |
| Ephys25 |  |  | B-E | A,C | B,C | H,I |  |
| Ephys26 |  |  | B-E | A-D | B,C | H,I |  |
| Ephys27 |  |  | B-E | A-D | B,C | H,I |  |
| Ephys28 |  |  | B-E | A,C | B,C | H,I |  |
| Ephys29 |  |  | B-E | A,C | B,C | H,I |  |
| Ephys30 |  |  | B-E | A,C | B,C | H,I |  |
| Ephys32 |  |  | B-E | A-D | B,C | H,I |  |
| Ephys33 |  |  | B-E | A-D | B,C | H,I |  |
| Ephys34 |  |  | B-E | A,C | B,C | H,I |  |
| Ephys36 |  |  | B-E | A,C | B,C | H,I |  |
| Ephys40 |  |  | A-E | A,C | B,C | H,I | Unit 8 in 20230626_s1 is 2nd example in Fig. 2A-C, S3A |
| Ephys41 |  |  | B-E | A-D | B,C | H,I |  |
| Ephys42 |  |  | B-E | A,C | B,C | H,I |  |
| beh104 | C,F |  |  |  |  |  |  |
| beh101 | F |  |  |  |  |  |  |
| beh102 | D-F |  |  |  |  |  |  |
| beh103 | A,B,F |  |  |  |  |  |  |
| Ephys72 |  |  |  |  |  | D,E | Unit 2 in 20240307_s1 is 1st example in Fig. S6E |
| Ephys73 |  |  |  |  |  | C,E | Unit 14 in 20240311_s1 is 2nd example in Fig. S6E |
| Ephys75 |  |  | B-E | A-D | B,C | H,I |  |
| Ephys76 |  |  | B-E | A,C | B,C | H,I |  |
| Ephys77 |  |  | B-E | A,C | B,C | H,I |  |
| Ephys78 |  |  | B-E | A,C | B,C | H,I |  |
| Ephys79 |  |  | B-E | A-D | B,C | H,I |  |
| Ephys81 |  |  | A-E | A,C | B,C | H,I | Unit 10 in 20240703_s1 is 4th example in Fig. 2A-C |
| Ephys83 |  |  | F-I |  |  | H,I |  |
| Ephys85 |  |  | F-I |  |  | H,I |  |
| Ephys87 |  |  | B-E | A,C | B,C | H,I |  |
| Ephys88 |  |  | A-E | A,C | A-C | H,I | Unit 5 in 20240905_s1 is 3rd example in Fig. 2A-C, S3A; 2nd in S5A |
| Ephys89 |  |  | B-E | A,C | B,C | H,I |  |
| Ephys92 |  |  | B-E | A-D | B,C | H,I |  |
| Ephys94 |  |  | F-I |  |  | H,I |  |
| Ephys96 |  |  | F-I |  |  | H,I |  |
| Ephys97 |  |  | F-I |  |  | H,I |  |
| Ephys100 |  |  | F-I |  |  | H,I |  |
| Ephys101 |  |  | B-E | A-D | B,C | H,I |  |
| Ephys102 |  |  | F-I |  | B,C | H,I |  |
| Ephys105 |  |  | B-E | A-E | B,C | H,I |  |
| Ephys106 |  |  | A-E | A-D | B,C | H,I | Unit 7 in 20240924_s1 is 6th example in Fig. 1A-C, S3A |
| Ephys107 |  |  | B-E | A,C,E | B,C | H,I | Unit 1 in 20240925_1 is 2nd example in Fig. S4E |
| Ephys108 |  |  | B-E | A,C | B,C | H,I |  |
| Ephys114 |  |  | F-I |  |  | H,I |  |
| Ephys119 |  |  | F-I |  |  | G-I |  |
| Ephys121 |  |  | F-I |  |  | H,I |  |
| Ephys122 |  |  | F-I |  |  | H,I |  |
| Ephys126 |  |  | B-E | A-D | B,C | H,I |  |
| Ephys127 |  |  | B-E | A-D | B,C | H,I |  |
| Ephys128 |  |  | B-E | A-D | B,C | H,I |  |

**Table S2.** Population Decoder Accuracy.

| | Decoder Accuracy Mean ± SEM (%) | | | | P-Value |
| --- | --- | --- | --- | --- | --- |
| Distance (mm) | Proximity | Map | Untuned | Shuffled | Proximity vs Map |
| **7** | 87.84 ± 1.210 | 81.14 ± 0.239 | 32.30 ± 1.767 | 11.17 ± 0.979 | 2.06 $\times 10^{-4}$, * |
| **9** | 78.87 ± 1.484 | 74.68 ± 0.330 | 15.89 ± 0.946 | 10.85 ± 0.738 | 3.30 $\times 10^{-4}$, * |
| **11** | 64.14 ± 2.238 | 82.57 ± 0.237 | 13.41 ± 1.045 | 11.42 ± 1.016 | 2.06 $\times 10^{-4}$, * |
| **13** | 50.60 ± 3.327 | 83.90 ± 0.213 | 13.04 ± 0.567 | 10.62 ± 1.008 | 2.06 $\times 10^{-4}$, * |
| **15** | 36.29 ± 1.824 | 82.50 ± 0.148 | 12.27 ± 0.789 | 10.78 ± 0.669 | 2.06 $\times 10^{-4}$, * |
| **17** | 30.73 ± 1.372 | 74.29 ± 0.377 | 12.01 ± 0.554 | 10.95 ± 0.690 | 2.06 $\times 10^{-4}$, * |
| **19** | 28.37 ± 1.284 | 63.16 ± 0.408 | 11.81 ± 0.384 | 10.69 ± 0.954 | 2.06 $\times 10^{-4}$, * |
| **21** | 28.91 ± 1.108 | 49.38 ± 0.439 | 11.63 ± 0.314 | 10.54 ± 0.828 | 2.06 $\times 10^{-4}$, * |
| **23** | 34.01 ± 0.670 | 52.12 ± 0.344 | 12.18 ± 1.011 | 11.09 ± 1.111 | 2.06 $\times 10^{-4}$, * |

\*: significant ( $p < 0.05$ ); n.s.: not significant

**Table S3.**
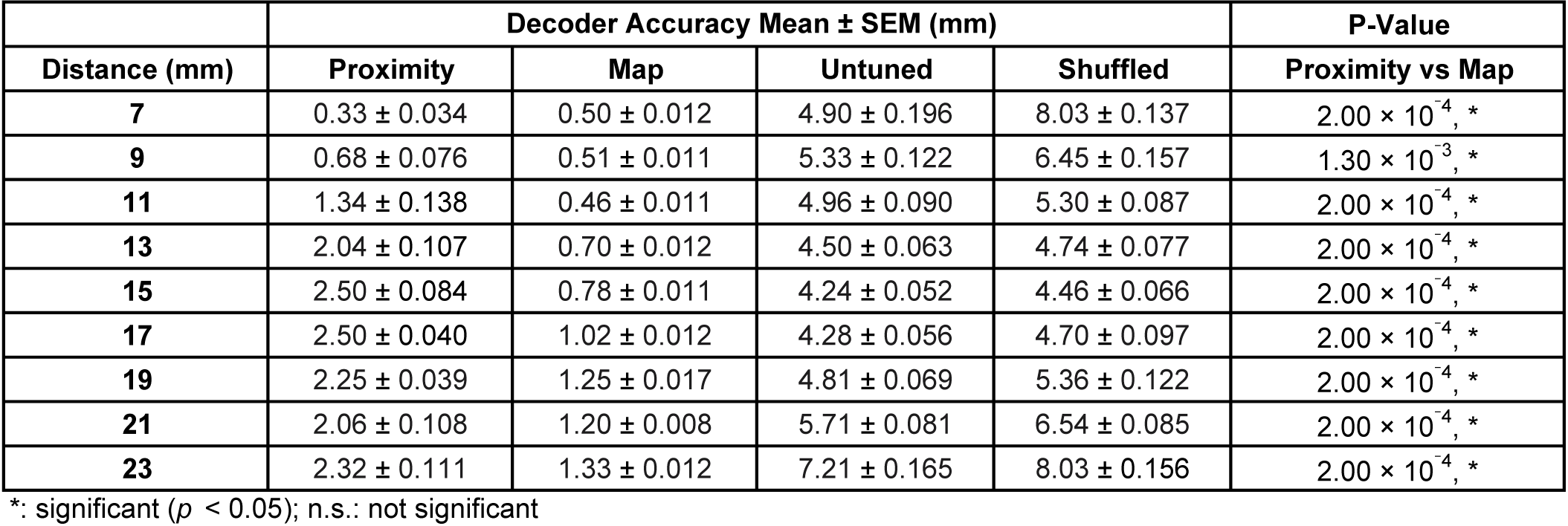
Population Decoder Error.

**Table S4.** Single Neuron Decoder Accuracy.

| Distance (mm) | Decoder Accuracy Mean ± SEM (%) | | P-Value |
| --- | --- | --- | --- |
|  | Proximity | Map | Proximity vs Map |
| **7** | 65.74 ± 1.319 | 59.09 ± 1.889 | 0.02, * |
| **9** | 40.83 ± 1.662 | 39.26 ± 2.253 | 0.73, n.s. |
| **11** | 31.44 ± 1.447 | 37.95 ± 2.203 | 0.02, * |
| **13** | 24.50 ± 1.312 | 37.34 ± 2.387 | 0, * |
| **15** | 18.75 ± 1.099 | 34.72 ± 2.277 | 0, * |
| **17** | 20.27 ± 1.195 | 27.63 ± 1.886 | 0, * |
| **19** | 20.68 ± 1.143 | 23.54 ± 1.824 | 0.15, n.s. |
| **21** | 20.69 ± 1.216 | 23.68 ± 1.722 | 0.05, n.s. |
| **23** | 26.44 ± 1.505 | 31.55 ± 2.313 | 0.03, * |

\*: significant ( $p < 0.05$ ); n.s.: not significant

**Table S5.** Single Neuron Decoder Error.

| | Decoder Accuracy Mean ± SEM (mm) | | P-Value |
| --- | --- | --- | --- |
| Distance (mm) | Proximity | Map | Proximity vs Map |
| 7 | 2.11 ± 0.115 | 2.94 ± 0.198 | 0, * |
| 9 | 3.34 ± 0.144 | 2.78 ± 0.166 | 0.21, n.s. |
| 11 | 3.89 ± 0.127 | 2.89 ± 0.157 | 0, * |
| 13 | 4.02 ± 0.101 | 2.85 ± 0.145 | 0, * |
| 15 | 3.76 ± 0.076 | 3.02 ± 0.146 | 0, * |
| 17 | 3.42 ± 0.072 | 3.32 ± 0.141 | 0.21, n.s. |
| 19 | 3.31 ± 0.077 | 3.45 ± 0.160 | 0.84, n.s. |
| 21 | 3.71 ± 0.103 | 3.76 ± 0.225 | 0.32, n.s. |
| 23 | 4.63 ± 0.143 | 4.58 ± 0.274 | 0.46, n.s. |

\*: significant ( $p < 0.05$ ); n.s.: not significant

**Table S6.** Statistics.

| Fig. | Comparison | Groups Tested | Measure (Mean ± SEM) | Statistical Test | Test Statistic | P-Value | Significance | DoF |
| --- | --- | --- | --- | --- | --- | --- | --- | --- |
| 3D | Population decoding accuracy | Proximity vs Map | see Table S2 | Wilcoxon rank-sum test | N/A | see Table S2 | see Table S2 | N/A |
| 3F | Population decoding error | Proximity vs Map | see Table S3 | Wilcoxon rank-sum test | N/A | see Table S3 | see Table S3 | N/A |
| 4B | Firing rates during wall passes | Full array vs. One whisker | 12.37 ± 2.986 vs. 8.61 ± 2.487 sp/s | Wilcoxon signed-rank | N/A | 1.27 $\times 10^{-9}$ | * | N/A |
| 5D | Preferred distance CDFs | Wild-type vs. TeLC populations | N/A | One-tailed K-S | N/A | 8.02 $\times 10^{-5}$ | * | N/A |
| 5E | Proportion of map units | Wild-type vs. TeLC populations | 10.34% vs. 3.90% | One-tailed Z-test | Z = 4.64 | 1.80 $\times 10^{-6}$ | * | N/A |
| 5E | Proportion of proximity units | Wild-type vs. TeLC populations | 23.31% vs. 32.09% | One-tailed Z-test | Z = -3.93 | 4.27 $\times 10^{-5}$ | * | N/A |
| 5E | Proportion of untuned units | Wild-type vs. TeLC populations | 48.20% vs. 49.92% | Two-tailed Z-test | Z = -0.68 | 0.5 | n.s. | N/A |
| 5E | Proportion of suppressed units | Wild-type vs. TeLC populations | 10.43% vs. 7.30% | One-tailed Z-test | Z = 2.12 | 0.02 | * | N/A |
| 5E | Proportion of ambiguous units | Wild-type vs. TeLC populations | 7.71% vs. 6.79% | Two-tailed Z-test | Z = 0.69 | 0.49 | n.s. | N/A |
| S1F, Left | C1 whisker midpoint | Awake vs. Anesthetized | -8.52 ± 0.633° vs. -5.88 ± 0.910° | Two-tailed paired t-test | t = -3.86 | 0.03 | n.s. ($\alpha$ = 0.0167) | 3 |
| S1F, Middle | C2 whisker midpoint | Awake vs. Anesthetized | -6.81 ± 0.867° vs. -4.72 ± 0.927° | Two-tailed paired t-test | t = -3.57 | 0.04 | n.s. ($\alpha$ = 0.0167) | 3 |
| S1F, Right | Greek whisker midpoint | Awake vs. Anesthetized | -4.76 ± 1.041° vs. -2.63 ± 0.714° | Two-tailed paired t-test | t = -4.55 | 0.0199 | n.s. ($\alpha$ = 0.0167) | 3 |
| S4B, Left | Firing rates of wall-activated, distance-tuned units | Wall passes vs. Baseline | 16.44 ± 1.640 vs. 6.36 ± 0.704 sp/s | One-tailed K-S | N/A | 9.26 $\times 10^{-28}$ | * | N/A |
| S4B, Right | Firing rates of wall-suppressed, distance-tuned units | Wall passes vs. Baseline | 4.47 ± 1.000 vs. 6.19 ± 1.076 sp/s | One-tailed K-S | N/A | 3.06 $\times 10^{-4}$ | * | N/A |
| S4C | Untuned unit firing rates | Wall passes vs. Baseline | 6.65 ± 0.687 vs. 6.90 ± 0.682 sp/s | Two-tailed K-S | N/A | 0.74 | n.s. | N/A |
| S4G, Middle | Wall-tuning category (ambiguous unit) | Non-closest vs. Closest distance | 32.59 ± 4.253 vs. 26.53 ± 4.107 sp/s | One-tailed Wilcoxon signed-rank test | N/A | 0.09 | n.s. | N/A |
| S4G, Right | Wall-tuning category (map unit) | Non-closest vs. Closest distance | 51.04 ± 2.49 vs. 26.09 ± 1.033 sp/s | One-tailed Wilcoxon signed-rank test | N/A | 1.27 $\times 10^{-4}$ | * | N/A |
| S5B | Single-neuron decoder accuracy | Proximity vs Map | see Table S4 | Wilcoxon rank-sum test | N/A | see Table S4 | see Table S4 | N/A |
| S5C | Single-neuron decoder error | Proximity vs Map | see Table S5 | Wilcoxon rank-sum test | N/A | see Table S5 | see Table S5 | N/A |
| S6H, Left | Baseline firing rates (untuned units) | TeLC vs. Wild-type populations | 9.34 ± 1.330 vs. 6.90 ± 0.682 sp/s | One-tailed K-S | N/A | 7.63 $\times 10^{-5}$ | * ($\alpha$ = 0.0125) | N/A |
| S6H, Right | Wall-pass firing rates (untuned units) | TeLC vs. Wild-type populations | 9.04 ± 1.390 vs. 6.65 ± 0.687 sp/s | One-tailed K-S | N/A | 1.06 $\times 10^{-4}$ | * ($\alpha$ = 0.0125) | N/A |
| S6I, Left | Baseline firing rates (tuned units) | TeLC vs. Wild-type populations | 5.60 ± 0.845 vs. 5.46 ± 0.670 sp/s | One-tailed K-S | N/A | 0.14 | n.s. ($\alpha$ = 0.0125) | N/A |
| S6I, Right | Wall-pass firing rates (tuned units) | TeLC vs. Wild-type populations | 20.25 ± 2.263 vs. 16.44 ± 1.640 sp/s | One-tailed K-S | N/A | 1.64 $\times 10^{-3}$ | * ($\alpha$ = 0.0125) | N/A |
| N/A | Firing rates during baseline | Fullarray vs. One whisker | 7.08 ± 1.424 vs. 7.08 ± 1.418 sp/s | Wilcoxon signed-rank | N/A | 0.59 | n.s. | N/A |

